# Perturbations of the ZED1 Pseudokinase Activate Plant Immunity

**DOI:** 10.1101/531665

**Authors:** D. Patrick Bastedo, Madiha Khan, Alexandre Martel, Derek Seto, Inga Kireeva, Jianfeng Zhang, Wardah Masud, David Millar, Jee Yeon Lee, Amy Huei-Yi Lee, Yunchen Gong, André Santos-Severino, David S. Guttman, Darrell Desveaux

## Abstract

The *Pseudomonas syringae* acetyltransferase HopZ1a is delivered into host cells by the type III secretion system to promote bacterial growth. However, in the model plant host *Arabidopsis thaliana,* HopZ1a activity results in an effector-triggered immune response (ETI) that limits bacterial proliferation. HopZ1a-triggered immunity requires the nucleotide-binding, leucine-rich repeat domain (NLR) protein, ZAR1, and the pseudokinase, ZED1. Here we demonstrate that HopZ1a can acetylate members of a family of ‘receptor-like cytoplasmic kinases’ (RLCK family VII; also known as PBS1-like kinases, or PBLs) and promote their interaction with ZED1 and ZAR1 to form a ZAR1-ZED1-PBL ternary complex. Interactions between ZED1 and PBL kinases are determined by the pseudokinase features of ZED1, and mutants designed to restore ZED1 kinase motifs can (1) bind to PBLs, (2) recruit ZAR1, and (3) trigger ZAR1-dependent immunity *in planta*, all independently of HopZ1a. A ZED1 mutant that mimics acetylation by HopZ1a also triggers immunity *in planta,* providing evidence that effector-induced perturbations of ZED1 also activate ZAR1. Overall, our results suggest that interactions between these two RLCK families are promoted by perturbations of structural features that distinguish active from inactive kinase domain conformations. We propose that effector-induced interactions between ZED1/ZRK pseudokinases (RLCK family XII) and PBL kinases (RLCK family VII) provide a sensitive mechanism for detecting perturbations of either kinase family to activate ZAR1-mediated ETI.

**AUTHOR SUMMARY:** All plants must ward off potentially infectious microbes, and those grown in large-scale crop operations are especially vulnerable to the rapid spread of disease by successful pathogens. Although many bacteria and fungi can supress plant immune responses by producing specialized virulence proteins called ‘effectors’, these effectors can also trigger immune responses that render plants resistant to infection. We studied the molecular mechanisms underlying one such effector-triggered immune response elicited by the bacterial effector HopZ1a in the model plant host *Arabidopsis thaliana*. We have shown that HopZ1a promotes binding between a ZED1, a ‘pseudokinase’ required for HopZ1a-triggered immunity, and several ‘true kinases’ (known as PBLs) that are likely targets of HopZ1a activity *in planta*. HopZ1a-induced ZED1-PBL interactions also recruit ZAR1, an *Arabidopsis* ‘resistance protein’ previously implicated in HopZ1a-triggered immunity. Importantly, ZED1 mutants that restore degenerate kinase motifs can bridge interactions between PBLs and ZAR1 (independently of HopZ1a) and trigger immunity *in planta*. Our results suggest that equilibria between active and inactive kinase domain conformations regulate ZED1-PBL interactions and formation of ternary complexes with ZAR1. Improved models describing molecular interactions between immunity determinants, effectors and effector targets will inform efforts to exploit natural diversity for development of crops with enhanced disease resistance.

## INTRODUCTION

Both plants and animals use cell membrane-spanning receptor kinases to sense extracellular molecular patterns produced by invading pathogens. While agonists of these receptors can trigger signal cascades that result in protective host immune responses (pattern-triggered immunity, or PTI), Gram-negative bacteria equipped with a type III secretion system (T3SS) can dampen such basal immune responses by delivery of effector proteins directly into host cells through the needle-like T3SS pilus. In plant cells, such T3SS-delivered effectors (T3Es) can be recognized by nucleotide-binding leucine-rich repeat (NLR) proteins, resulting in ‘effector-triggered immunity’ (ETI), which is often accompanied by a hypersensitive cell death response (HR) [1]. NLRs are members of the STAND (signal transduction ATPases with numerous domains) class of P-loop NTPases that also includes the NOD-like receptors of animal cells [2]. These proteins share common central nucleotide-binding and carboxy-terminal LRR domains, but are preceded by distinct classes of amino-terminal domains – plant NLRs have either ‘coiled-coil’ (CC) or ‘Toll/Interleukin-1 Receptor’ (TIR) domains at their amino-termini [2–5].

Plant NLRs become activated either by direct sensing of effectors, or by indirect sensing of effector-modified substrates [6]. Such effector-modified substrates can be virulence targets whose functions limit bacterial growth, or mimics of these virulence targets referred to as ‘decoys’ [7]. For example, the *Arabidopsis* kinase PBS1 (Avr<u>P</u>ph<u>B</u>-<u>S</u>usceptible <u>1</u>), is cleaved by the *P. syringae* T3E HopAR1 (previously known as AvrPphB), resulting in activation of the *Arabidopsis* NLR RPS5 [8,9]. There are 45 PBS1-like kinases (PBL kinases, or PBLs) that together with PBS1 comprise receptor-like cytoplasmic kinase (RLCK) subfamily VII. At least eight of these additional PBLs are also sensitive to cleavage by HopAR1, including BIK1 and PBL1, which are both involved in plant immunity [10]. PBS1 therefore appears to function as a decoy that mimics HopAR1 virulence targets to activate RPS5-mediated ETI [10].

The *Arabidopsis* NLR ZAR1 (Hop<u>Z</u>-<u>A</u>ctivated <u>R</u>esistance <u>1</u>) belongs to the coiled-coil class of plant NLRs and is required for ETI against at least three distinct T3Es: HopZ1a and HopF2a from *Pseudomonas syringae*, and AvrAC from *Xanthomonas campestris* [11–13]. In addition to ZAR1, recognition of all three T3Es requires members of RLCK family XII. Members of this RLCK family lack at least one of several well-established and highly conserved kinase motifs, and are therefore considered to encode pseudokinases [14–16]. Recognition of the acetyltransferase HopZ1a requires the RLCK XII pseudokinase ZED1 (Hop<u>Z E</u>TI-<u>D</u>eficient <u>1</u>), which is encoded in a gene cluster with seven additional ZED1-related pseudokinases (ZRKs). ZRK1/RKS1 (hereafter ZRK1) and ZRK3 pseudokinases are required for recognition of AvrAC and HopF2a, respectively [12–14]. ZED1, ZRK1 and ZRK3 have all been shown to interact with ZAR1 in planta, although different roles in T3E recognition have been proposed [13,15,16]. ZED1 is acetylated by HopZ1a, but a *ZED1* knockout does not appear to influence the virulence of *P. syringae* [14]. As such, it has been proposed that ZED1 is a decoy substrate monitored by ZAR1 to guard against the acetylation of other (kinase) substrates of HopZ1a [14–16]. In contrast, ZRK1 functions as an adaptor for ZAR1 by recruiting PBL2 (<u>PB</u>S1-like kinase 2) proteins uridylated by AvrAC to elicit ETI [12,17,18]. Recent structural studies indicate that binding of uridylated PBL2 to ZAR1-bound ZRK1 promotes ADP/ATP exchange by ZAR1 and results in formation of a pentameric ‘resistosome’ similar to the inflammasomes and apoptosomes formed by mammalian NLRs [17,18]. ZRK3 is also thought to act as a ZAR1 adaptor for an as-yet-unidentified kinase that is ADP-ribosylated by the *P. syringae* T3E HopF2a [13]. Overall, the ZED1/ZRK family of pseudokinases have expanded the T3E recognition capacity of ZAR1 by directly sensing T3E modifications and/or serving as adaptors that bridge interactions between the NLR and T3E-modfied plant kinases.

Highly-conserved kinase motifs were initially identified by comparative sequence analysis [19,20], but since then, determination of the molecular structures of an ever-increasing number of diverse kinase domains in the presence or absence of nucleotides, metal ions, substrate peptides, and/or pharmacological inhibitors has provided rational explanations for the functional importance of these motifs [21]. The kinase domain nucleotide-binding pocket is formed by a cleft between a mostly β-stranded amino-terminal lobe and an α-helical carboxy-terminal lobe. In the amino-terminal lobe, a glycine-rich motif in the loop connecting strands β1 and β2 (‘G-loop’) acts as a lid through backbone interactions with the β- and γ-phosphates of bound ATP [22]. Appropriate positioning of ATP phosphates also requires metal ions (usually Mg^2+^ or Mn^2+^), a lysine from strand β3 that is stabilized by a salt-bridge with a glutamate from helix αC, and an aspartate from the ‘DFG motif’. The DFG motif also defines the beginning of the ‘activation loop’, a conformationally flexible (and often disordered) region of 20-35 amino acids that typically contains serine and/or threonine residue that can be phosphorylated (by autophosphorylation or by upstream kinases). Phosphorylation of the activation loop can stabilize an open conformation (i.e. a ‘DFG-in’ orientation, and a fully-assembled hydrophobic ‘regulatory spine’) that is associated with active kinases [23]. This open conformation allows substrate peptides to approach the γ-phosphate (when appropriately-positioned, as described above) for phospho-transfer by the aspartate from an ‘HRD motif’ in the catalytic segment. These dynamic structural features play essential roles in regulating kinase activity as well as modulating their interactions with other proteins [21,24]. ZED1 and the ZRKs have degeneracies in nearly all of these canonical kinase motifs, and as such they may all represent pseudokinases that perform T3E sensing functions similar to those proposed for ZED1, ZRK1 and ZRK3 [12–14,16].

Early surveys of the complete kinase domain complements from organisms including human, mouse, *Caenorhabditis elegans*, *Dictyostelium* and *Arabidopsis* found that pseudokinases like ZED1 account for ∼10% of the kinase domains in higher eukaryotes and 20% of *Arabidopsis* RLCKs [25,26]. Plant pseudokinases are emerging as important mediators of development and immunity [27–29]. Rather than mere non-functional kinases resulting from relaxed (or absent) selective constraints, there is instead a growing body of evidence indicating that pseudokinases are important signaling molecules in spite of their predicted lack of catalytic activity, often functioning as adaptors or scaffolds that modulate the activity of true kinase partners [25,28,30,31]. In this report we show that the *P. syringae* T3E HopZ1a induces interactions between ZED1 and PBL kinases and promotes the formation of a ZAR1-ZED1-PBL ternary complex. The pseudokinase features of ZED1 mediate its conditional interactions with PBL kinases, recruitment of ZAR1, and importantly, its regulation of ZAR1-mediated immunity. Overall, our results support the hypothesis that ZED1-PBL (and more generally ZRK-PBL) pseudokinase-kinase interplay provides a sensor for perturbations of host kinases by pathogen-delivered effector proteins.

## RESULTS

### HopZ1a binds and acetylates PBL kinases

In order to assess whether PBL kinases are plausible targets of HopZ1a, we tested whether HopZ1a can interact with members of this family. Using a yeast two-hybrid (Y2H) screen (S1 Fig; interaction scheme A) with all 46 PBL kinases as prey and HopZ1a as bait (wild-type or a C216A catalytic site mutant) we found strong binding between nine PBL kinases (PBL21, PBL27, PBL8, PBL2, PBL3, PBL4, PBL18, PBL15 and PBL13) and HopZ1a^C216A^, but no interactions were observed with the wild-type effector, HopZ1a^wt^ (Fig 1A).

**Fig 1.**
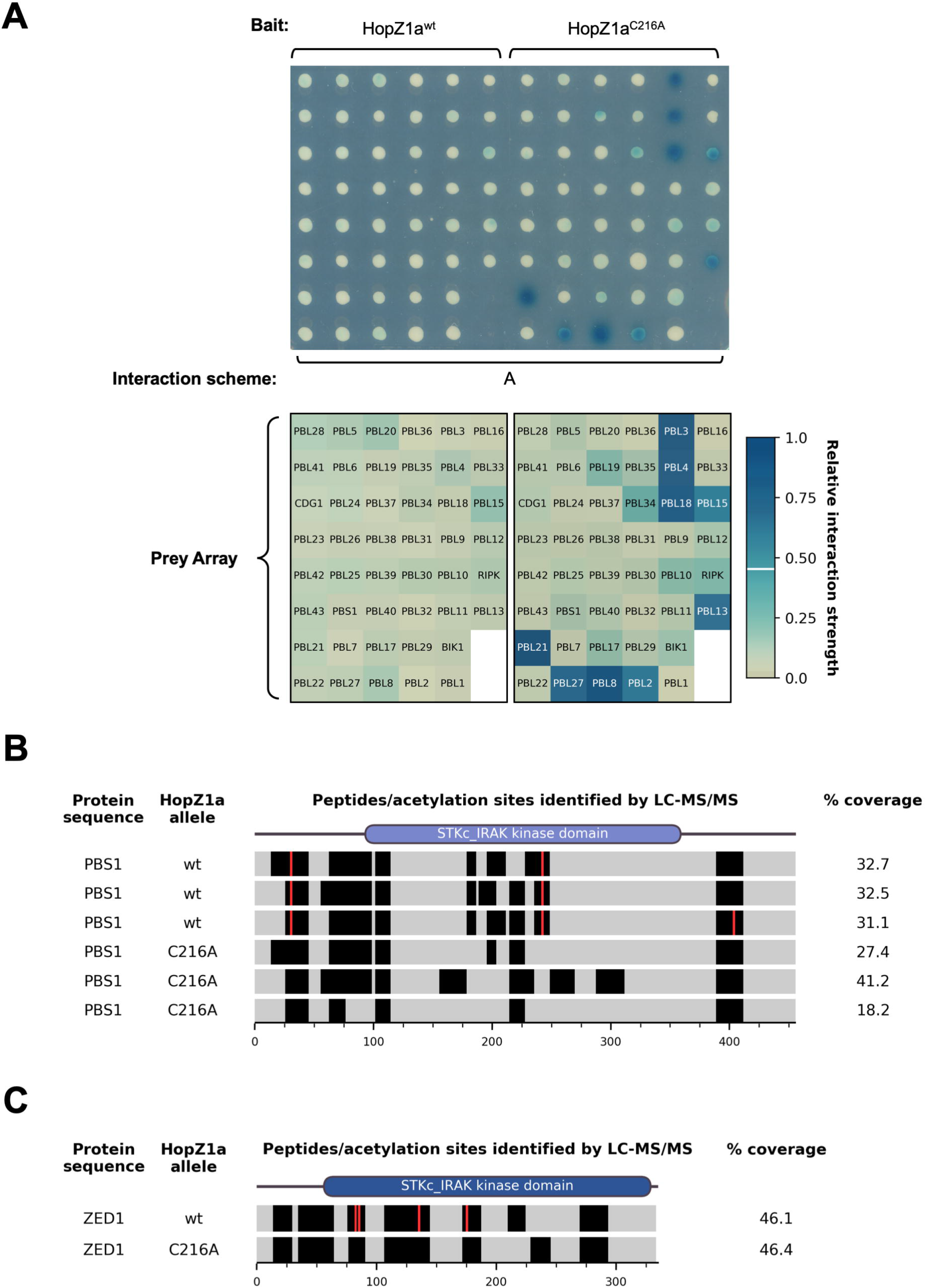
HopZ1a binds to PBL kinases and acetylates PBS1. (A) Yeast two-hybrid interaction assay using wild-type or catalytically-inactive HopZ1a alleles as bait and an array of 46 PBL kinases as prey. Yeast colonies on an X-Gal reporter plate are shown on top. A grid describing the PBL array layout is shown below each half of the plate, with the colours of array positions representing the averaged colony colour for each bait-prey interaction. We used a ‘relative interaction strength’ metric derived from these averaged colony colours to discriminate strong interactions from the background (see S2 Fig; Materials and Methods); prey array positions with interaction strength ≥ 0.455 (i.e. strong interactions) were assigned white labels while other positions were assigned black labels). The interaction strength threshold is indicated in the colour-bar at right with a horizontal white line. Interaction scheme refers to S1 Fig. (B) LC-MS/MS identification of PBS1 residues acetylated by HopZ1a. The PBS1 protein sequence is indicated as horizontal bars for three experimental replicates in the presence of each HopZ1a allele (wild-type HopZ1a or the catalytically-inactive mutant, HopZ1a^C216A^. Black bands indicate peptides that were reliably detected by the mass-spectrometry instruments, while peptides not detected with high confidence are indicated by grey shading. (C) LC-MS/MS identification of ZED1 residues acetylated by HopZ1a. Peptide detection and acetylation sites are presented as for panel B above. Note that in addition to S84, T87 and T177, S137 was also acetylated, although this modification was not consistently observed across multiple independent experiments.

We hypothesized that failure to observe binding between HopZ1a^wt^ and PBLs may reflect efficient enzymatic activity and rapid substrate turnover (i.e. ‘catch and release’ catalysis). Such interactions would be stabilized in the case of HopZ1a^C216A^ due to frustrated enzymatic activity. To test whether HopZ1a can acetylate PBL kinases, we co-expressed FLAG-tagged derivatives of both HopZ1a and PBS1 in yeast, prepared anti-FLAG immunoprecipitates from yeast cell extracts and subjected these to liquid chromatography-tandem mass spectrometry (LC-MS/MS) analysis. Although PBS1 was a weaker HopZ1a interactor, we chose to test its acetylation by HopZ1a based on its established role in ETI. We observed three PBS1 peptides with mass increases of 42 Da (corresponding to addition of an acetyl group) when PBS1 was co-expressed with HopZ1a^wt^ but not with HopZ1a^C216A^, indicating HopZ1a-mediated acetylation. Fragmentation analysis of these three peptides established that the specific sites acetylated were T32, S244, and S405 (Fig 1B; S3 Fig; S1 File). These results support our hypothesis that HopZ1a can acetylate PBL kinases.

### HopZ1a promotes binding between ZED1 pseudokinase and PBL kinases

Given the previously reported finding that AvrAC promotes interactions between PBL2 and ZRK1 [12], we sought to develop an assay to investigate whether HopZ1a can promote interactions between ZED1 and PBL kinases. We devised a yeast three-hybrid (Y3H) assay wherein interactions between a ZED1/ZRK bait protein and PBL preys were assessed in the absence and presence of single-copy chromosomal integrations of a T3E of interest (S1 Fig, interaction schemes A, B). As proof-of-principle we integrated *Xanthomonas* AvrAC at the yeast *ho* locus and investigated its influence on ZRK1 interactions with PBL kinases. In the absence of AvrAC expression, ZRK1 exhibited high-affinity binding to several PBLs (PBL21, PBL17, PBL8 and PBL15) and moderate binding to several others, including PBL2 (S4 Fig). Binding between ZRK1 and PBL2 was enhanced by the presence of AvrAC (as was binding between ZRK1 and PBL3 / PBL29; S4 Fig), indicating that our Y3H system can be used to examine modulation of ZRK/PBL interactions by T3Es.

We then integrated *hopZ1a* (wild-type or C216A alleles) at the yeast *ho* locus and examined its influence on ZED1-PBL interactions. Only weak ZED1-PBL interactions were observed in the absence of effector (Fig 2B; Fig 3B, column 1) or when HopZ1a^C216A^ was expressed (Fig 2A; Fig 3B, column 3). Notably, expression of wild-type HopZ1a resulted in enhanced interactions between ZED1 and 11 PBL kinases (PBL21, PBL22, PBL5, PBS1, PBL27, PBL17, PBL8, PBL4, PBL9, PBL15, and PBL13), with the strongest interactions occurring between ZED1 and PBL5 / PBL27 / PBL17 / PBL8 / PBL15 (Fig 2A; Fig 3B, column 2). These results indicate that HopZ1a activity can promote interactions between the ZED1 pseudokinase and PBL kinases.

**Fig 2.**
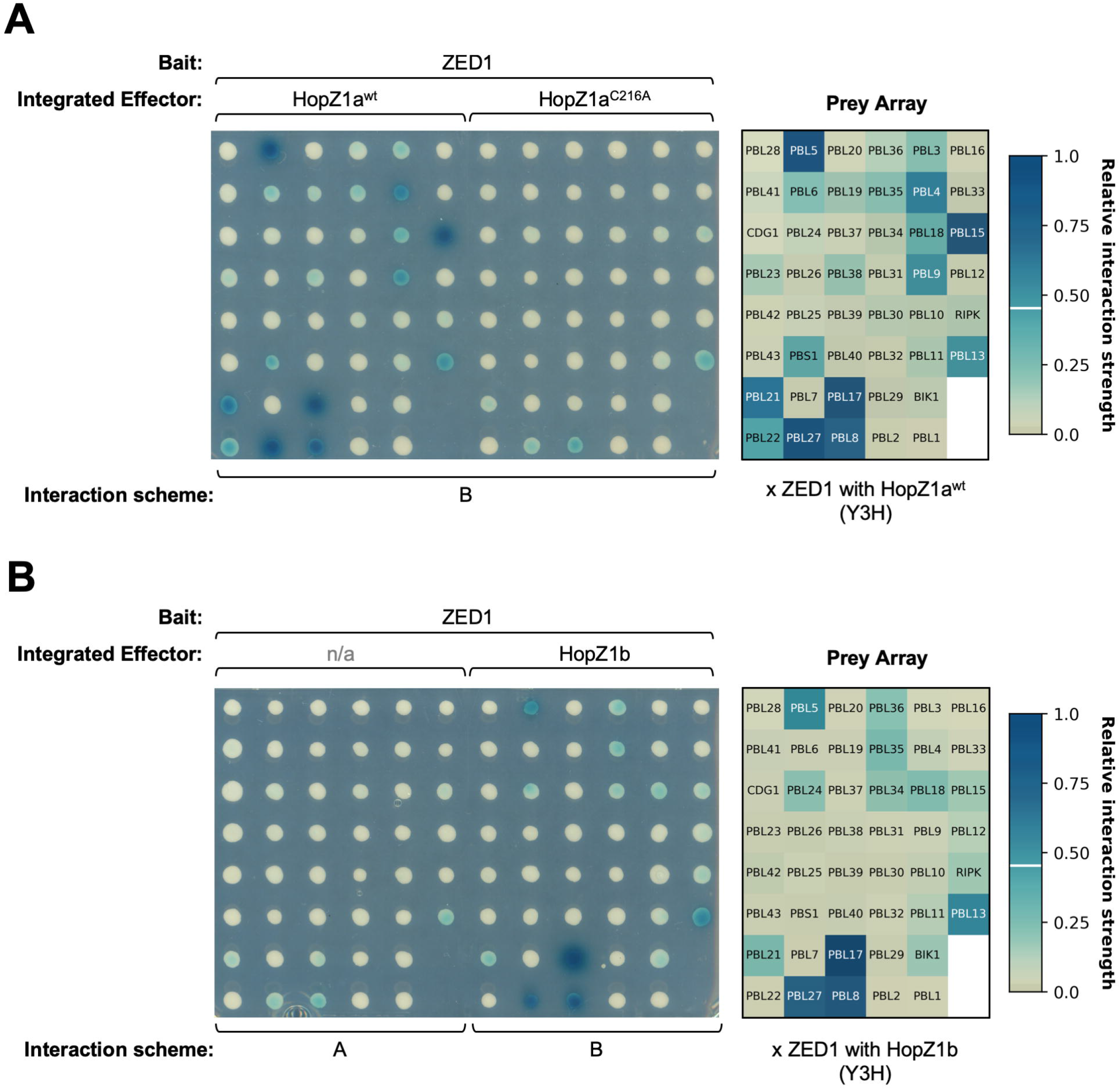
HopZ1a promotes binding between ZED1 and several PBL kinases. Y3H assays testing interactions between ZED1 and 46 PBLs in the presence of HopZ effector alleles are contrasted with a Y2H assay in the absence of effectors. (A) HopZ1a^wt^ (left) or HopZ1a^C216A^ (center) were expressed from the chromosome (see Materials and Methods) with ZED1 as bait and 46 individual PBLs as prey. The prey array layout shown at right represents PBL interactions with ZED1 in the presence of HopZ1a^wt^. The colours of the labels at each array position (white or black) are determined by the relative interaction strength (see S2 Fig and Materials and Methods). Interaction scheme refers to S1 Fig. (B) ZED1 bait / PBL prey interactions in the absence of chromosomally expressed effector (left) or in the presence of HopZ1b (center). The prey array layout shown at right represents PBL interactions with ZED1 in the presence of HopZ1b. Interaction schemes refer to S1 Fig.

**Fig 3.**
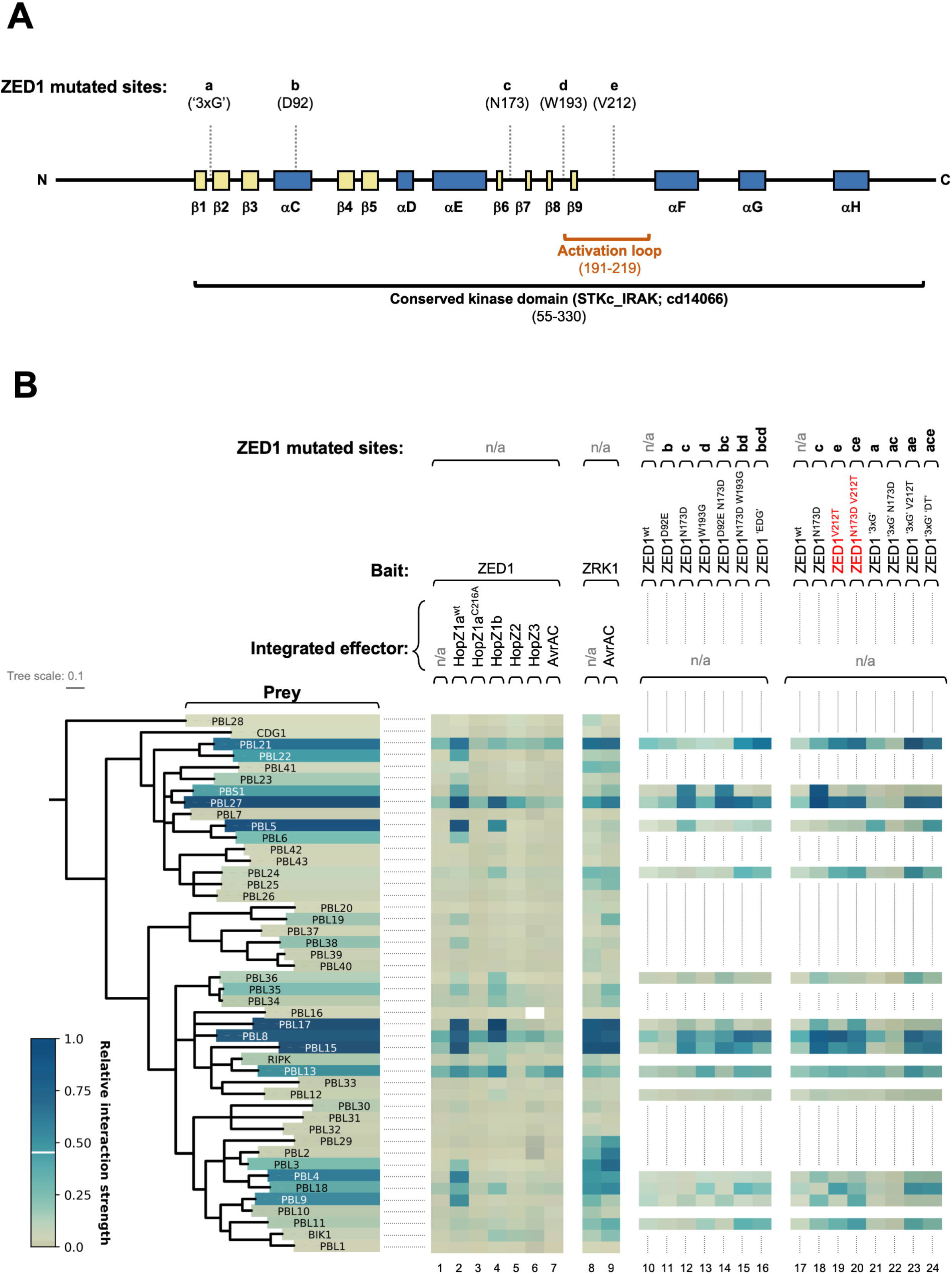
ZED1 mutagenesis targets, phylogenetic analysis of PBL kinase domains, and a summary of ZED1-PBL binding interactions. (A) A linear representation of the ZED1 sequence showing the positions of mutated sites (labeled a-e, according to their descriptions in the text), and predicted secondary structure elements are labeled according to homologous features present in the well-studied kinase, PKA [85–88]. β-strands are coloured yellow, while α-helices are coloured dark blue, as for the three-dimensional structural models of ZED1 presented in S7 Fig (panel A), S10 Fig (panel A) and S17 Fig. The activation loop sequence is highlighted with an orange bracket. (B) *Left* – phylogenetic tree showing relationships between the kinase domains of RLCK family VII (PBS1-like) kinases. Labels for tree leaves are coloured according to yeast colony colours from the HopZ1a^wt^-dependent interactions presented in Fig 2A. *Right* - condensed graphical summary of the Y2H/Y3H interaction data presented in Fig 2, S4 Fig, S5 Fig, S8 Fig and S9 Fig. Subsets of the five ZED1 mutagenesis targets (sites a-e) that are altered in a given ZED1 allele are indicated at the top of each column. Mutant ZED1 alleles whose expression causes induction of ETI *in planta* are highlighted with red text. The colour-bar at lower left depicts the range of relative interactions strengths, as defined in S2 Fig (see Materials and Methods).

In order to assess whether stimulation of ZED1-PBL binding was a property specific to HopZ1a, we also created yeast strains with chromosomal integrations of genes encoding both closely-related (HopZ1b; 65% amino acid identity to HopZ1a over 349 non-gapped sites) and more distantly-related HopZ family members (HopZ2, HopZ3; 26% and 23% identical to HopZ1a over 335 and 306 non-gapped sites, respectively). Interactions between ZED1 and PBLs were assessed in the presence of each of these HopZ family effectors or with the unrelated effector AvrAC (Fig 2B; S5 Fig; Fig 3B, columns 2-7). Like HopZ1a, HopZ1b promoted strong interactions between ZED1 and PBL27 / PBL17 / PBL8, but in contrast, interactions between ZED1 and PBL5 / PBL15 / PBL21 were weaker with HopZ1b than with HopZ1a, and HopZ1b did not promote interactions between ZED1 and PBL22 / PBS1 / PBL4 / PBL9 (Fig 2 and Fig 3B, columns 2, 4). Notably, the interactions between ZED1 and PBLs were unchanged by the expression of HopZ2, HopZ3, or AvrAC (Fig 3B, columns 5-7; S5 Fig, panels A, B) although integrated effectors are expressed to comparable levels (S5 Fig, panel C), demonstrating that promotion of ZED1/PBL interactions is specific to HopZ1a (and to a lesser extent, HopZ1b).

### Perturbation of ZED1 pseudokinase features modulates PBL interactions

The kinase domains of ZED1 and the ZRKs have diverged significantly from those of true kinases, with degeneracies in one or more of the established kinase motifs (S6 Fig) [14]. They also generally lack kinase activity under *in vitro* conditions that are sufficient for canonical kinases to phosphorylate model substrates [14,32]. To determine the importance of ZED1 pseudokinase features for (HopZ1a-induced) binding to PBLs, we identified conspicuous ZED1 sequence elements based on their divergence from canonical kinase motifs and designed mutations intended to ‘restore’ these degenerate motifs. We made a series of ZED1 variants with various combinations of mutations at five different sites (Fig 3A), described below.

**a)** The glycine-rich motif (‘GXGGFG’) between strands β1 and β2 is unrecognizable in ZED1 (S6 Fig, panel B). The triple mutant (S56G W58G F61G; hereafter referred to as ZED1^‘3xG’^) restores a glycine-rich motif at this position.
**b)** The salt-bridge formed between the lysine from strand β3 and the glutamate from helix αC is evident in the crystal structure of the *Arabidopsis* receptor kinase BRI1 (between K911 and E927; S7 Fig, panel B) [33], while in a homology-based structural model of ZED1 the sidechains of the corresponding residues (K76 and D92) are too far apart (S7 Fig, panel A). ZED1^D92E^ extends the acidic side chain at this position by a single CH_2_ group to restore an αC glutamate.
**c)** ZED1^N173D^ restores the active site aspartate of the HRD motif required for phospho-transfer in active kinases.
**d)** In ZED1 the DFG motif glycine is replaced by a tryptophan (S6 Fig, panel B; S7 Fig, panel A). ZED1^W193G^ restores the activation loop DFG motif.
**e)** ZED1 is unique, even among the ZRKs, in that its activation loop is completely devoid of potential phospho-accepting serine or threonine residues (S6 Fig, panel B). We thus created ZED1^V212T^, since this position is an absolutely conserved threonine in all of the other kinase sequences considered in our analysis, and its phosphorylated derivative has been observed in a crystal structure of the *Arabidopsis* BAK1 kinase domain [34].

We tested the PBL-binding activity of each of these rationally-designed ZED1 mutations (individually and in various combinations) both in the absence and presence of HopZ1a expression using Y2H and Y3H assays. We examined the binding of ZED1 mutants to a restricted subset of 15 PBLs that includes ten representatives with robust HopZ1a-induced binding affinity to wild-type ZED1 as well as five with relatively weak HopZ1a-dependent ZED1-binding activity (PBL24, PBL36, PBL18, PBL12, PBL11).

Our mutational analysis found that ZED1^N173D^ (HRD restoration), ZED1^W193G^ (DFG restoration) and ZED1^V212T^ (activation loop restoration) all gained affinity for several PBL kinases in the absence of HopZ1a, with ZED1^N173D^ and ZED1^V212T^ showing the strongest interactions (S8 Fig and S9 Fig; Fig 3B, compare columns 12, 13, 18, 19 with columns 1, 10, 17). These mutations induced HopZ1a-independent ZED1 binding to PBL kinases showing both strong (PBL21, PBL27, PBL15, PBL17, PBL4) and weak (PBL18) HopZ1a-dependent binding. HopZ1a-independent binding between PBS1 and ZED1 was only observed in the context of the ZED1^N173D^ mutation (i.e. HRD restoration; ZED1^N173D^ and ZED1^D92E N173D^; S8 Fig and S9 Fig; Fig 3B, columns 12, 14, 18). The triple mutant ZED1^‘3xG’^ (G-loop restoration) disrupted HopZ1a-induced PBL binding, and only moderate HopZ1a-independent binding was observed (S9 Fig, panel B; Fig 3B, column 21). Combining ZED1 mutations differentially influenced subsets of PBL interactions. For example, combining N173D with W193G (ZED1^N173D W193G^; S8 Fig, panel B; Fig 3B, column 15) or with D92E and W193G (ZED1^‘EDG’^; S8 Fig, panel B; Fig 3B, column 16) resulted in a loss of PBS1 binding (observed with ZED1^N173D^ and ZED1^D92E N173D^; S8 Fig; Fig 3B, columns 12, 14, 18) but increased ZED1 binding affinity for PBL21 and PBL24 (not observed with any of the three individual mutations; S8 Fig, panel A and S9 Fig, panel A; Fig 3B, columns 11, 12, 13, 18). When we combined N173D with the restored G-loop (ZED1^‘3xG’ N173D^), all binding to PBLs was abolished whether HopZ1a was present or not (S9 Fig, panel B; Fig 3B, column 22). However, a subset of interactions was restored in the context of a phospho-accepting activation loop in ZED1 (ZED1^‘3xG’ V212T^ and ZED1^‘3xG’ ‘DT’^, which adds both N173D and V212T; S9 Fig, panel B; Fig 3B, columns 23, 24). These results suggest that the effects of ZED1 mutations differentially affect binding affinity for distinct PBLs, indicating that PBLs differ in their abilities to interact with ZED1 through perturbation of (pseudo)kinase features either through mutation or by T3E modification.

### HopZ1a acetylation sites influence ZED1/PBL interactions

Based on the results of our ZED1 mutagenesis, we hypothesized that acetylation of one or more sites on ZED1 might influence ZED1-PBL binding by shifting between ‘pseudokinase-like’ and ‘kinase-like’ states. We have previously described acetylation of ZED1 by HopZ1a at T125 and T177 [14], and subsequent experiments have indicated that S84 and T87 can also be acetylated by HopZ1a (Fig 1C; S3 Fig; S1 File). We therefore made glutamine substitutions as potential mimics of acetylated serine and/or threonine residues based on similarities in sidechain length and branching structure (Fig 4A). We tested the PBL-binding capacity of three ZED1 glutamine substitution mutants: ZED1^T177Q^, which is proximal to the (non-catalytic) active site residue N173, as well as ZED1^S84Q^ and ZED1^T87Q^, which are both predicted to be part of a solvent-exposed surface of helix αC (S7 Fig, panel A; S10 Fig, panel A). None of the ZED1 glutamine substitution mutants (or any of the three possible pairwise combinations) resulted in HopZ1a-independent binding to any of the PBLs tested (S10 Fig, panel B). In contrast, each of the mutants bearing the T177Q substitution (ZED1^T177Q^, ZED1^S84Q T177Q^, ZED1^T87Q T177Q^) lost all capacity for HopZ1a-dependent binding (S10 Fig, panel B), indicating that ZED1^T177Q^ represents a loss-of-function allele.

**Fig 4.**
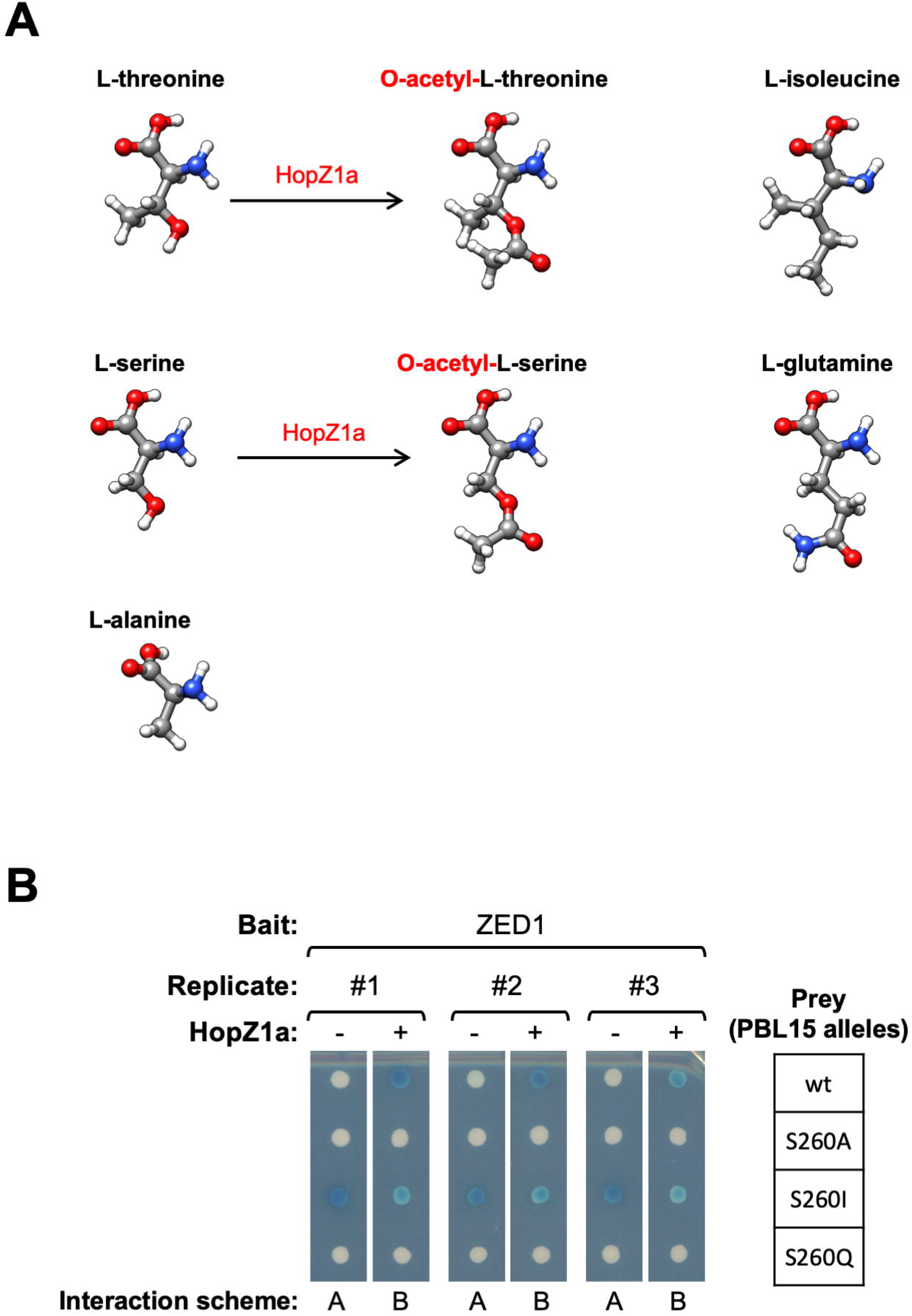
Isoleucine substitution of a PBL activation loop phospho-accepting residue mimics acetylation by HopZ1a. (A) Comparison of the molecular structures of phospho-accepting residues L-threonine and L-serine, their acetylated derivatives, candidate acetyl-mimetic residues L-isoleucine and L-glutamine, and L-alanine, a presumed loss-of-function substitution that should block both phosphorylation and acetylation. (B) Mutant alleles of PBL15 were tested in both Y2H (ZED1 bait / PBL15 prey) and Y3H contexts (ZED1 bait / PBL15 prey with HopZ1a expression) to screen for substitutions that confer HopZ1a-independent ZED1 binding activity. Interaction schemes refer to S1 Fig.

We also tested whether the PBS1 acetylation sites described above (Fig 1B; S3 Fig; S1 File) are important for its interaction with ZED1. Similar to our ZED1 findings, we found that none of the PBS1 glutamine substitutions (alone or in combination) enable HopZ1a-independent ZED1-binding activity (S10 Fig, panel C). Furthermore, as for ZED1 T177, each of the PBS1 mutants bearing a glutamine substitution of S244 lost capacity for HopZ1a-dependent binding to ZED1 (S10 Fig, panel C). The combined insights from our mutagenesis of ZED1 and PBS1 acetylation sites suggest that glutamine does not mimic acetylated serine/threonine residues and may instead act as a loss-of-function mutation by preventing acetylation and/or phosphorylation.

We also considered isoleucine substitutions as possible mimics of acetylated serine and/or threonine residues based on similarities in hydrophobicity and sidechain branching (Fig 4A). We therefore tested glutamine and isoleucine (possible acetyl-mimics) as well as alanine (loss-of-function) substitutions in PBL15, which was selected as a representative PBL based on its strong HopZ1a-dependent binding to ZED1 (stronger than PBS1; Fig 2A). We mutated PBL15 S260, an activation loop residue equivalent to PBS1 S244 (which is acetylated by HopZ1a; Fig 1B). Of these three substitutions of PBL15 S260, only PBL15^S260I^ demonstrated ZED1 binding in the absence of HopZ1a activity (Fig 4B). The strength of this interaction was comparable to the ZED1-PBL15^wt^ binding induced by HopZ1a, and HopZ1a-induced ZED1 binding was abolished by the alanine and glutamine substitutions of S260. Overall, these results suggest that isoleucine substitutions can mimic acetylation of serine (and possibly threonine) residues and that acetylation of the activation loops of PBL kinases by HopZ1a is sufficient to promote their interactions with ZED1.

### HopZ1a promotes the formation of a ZAR1-ZED1-PBL ternary complex

Having established that HopZ1a is able to stimulate protein-protein interactions between ZED1 and PBLs, we wished to investigate whether HopZ1a can also promote formation of a ternary complex between PBLs, ZED1, and ZAR1. For this purpose, we developed a yeast four-hybrid system where ZAR1 baits and PBL kinase preys (11 with HopZ1a-dependent ZED1 binding) were expressed in the presence or absence of chromosomally-integrated *hopZ1a* (wild-type or the C216A catalytic site mutant) and *ZED1* (S1 Fig, interaction scheme C). Since modulation of inter-domain contacts is known to be important for the activation of plant NLRs [17,18,35,36], we created various domain truncations of ZAR1 (panel A in both Fig 5 and S11 Fig) in order to investigate domain-specific ZAR1 binding interactions with ZED1/PBLs that might be modulated by HopZ1a activity. Full-length ZAR1 bait (ZAR1^wt^) interacted with prey PBL kinases (PBL5, and weaker interactions with PBL17, 4, 18, 15) when co-expressed with both ZED1 and HopZ1a^wt^, but not when co-expressed with ZED1 and HopZ1a^C216A^ (Fig 5B, compare columns 4 and 6). These interactions were dependent on ZED1 expression (Fig 5B, compare columns 3 and 4) and also required the ZAR1 LRR domain, since no interactions were observed with the ZAR1^ΔLRR^ construct (Fig 5B, compare columns 4 and 12). Consistent with this finding, the isolated LRR domain displayed weak HopZ1a^wt^/ZED1-dependent PBL interactions, but the CC and NB domains did not (S11 Fig, panel B, columns 4, 10, 16). Interestingly, ZAR1^ΔCC^ showed stronger HopZ1a/ZED1-dependent interactions with 9 of the 11 PBLs (Fig 5B, compare columns 4 and 20), suggesting that the ZAR1 coiled-coil domain negatively regulates ZAR1-ZED1-PBL complex formation.

**Fig 5.**
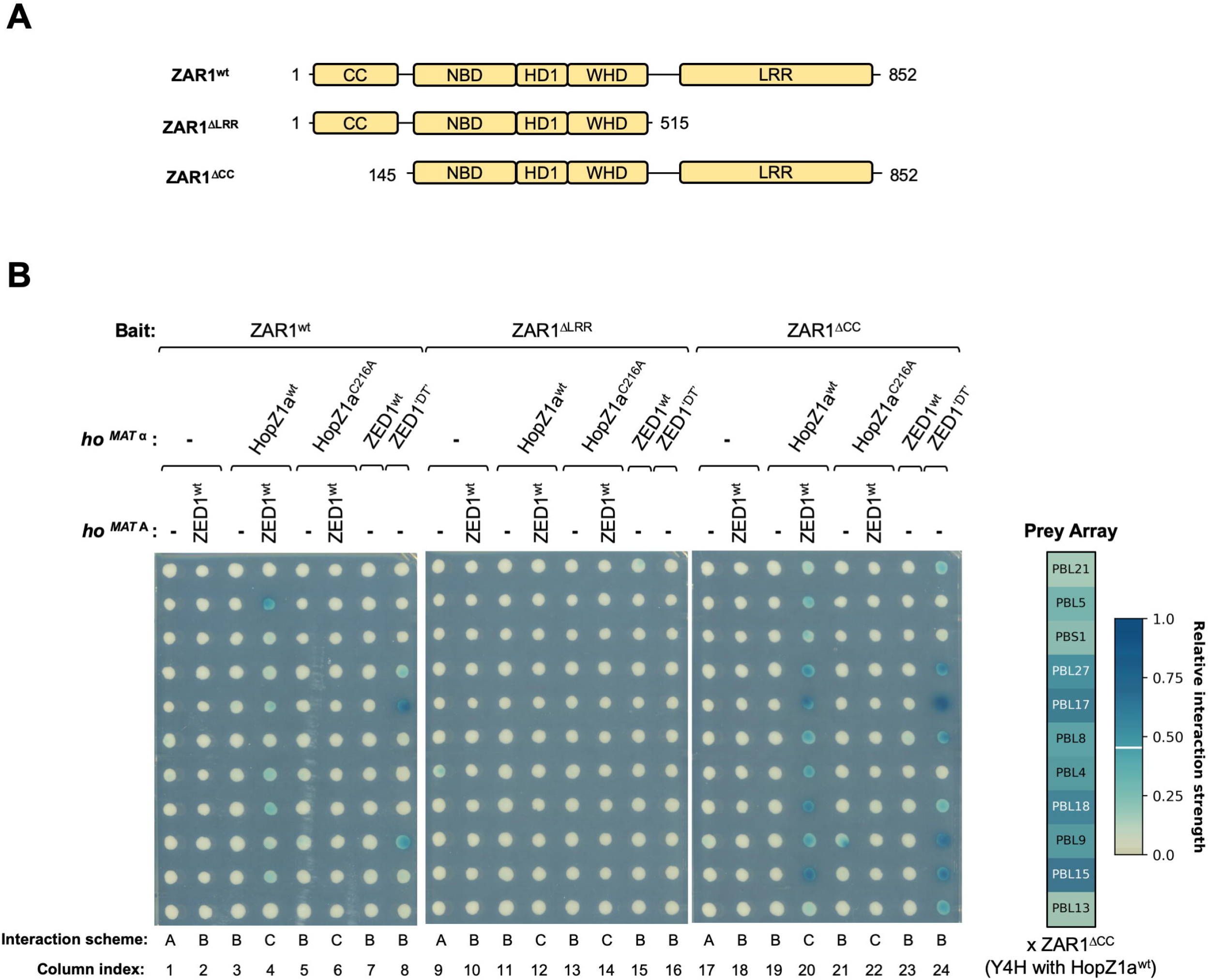
ZAR1-ZED1-PBL ternary complexes are formed in the presence either of HopZ1a activity or of a mutant ZED1 allele. (A) Schematic showing ZAR1 domain truncation boundaries. Subdomains of the central nucleotide-binding region are labeled as described by Wang et al [17,18]: NBD, nucleotide-binding domain; HD1, helical domain 1; and WHD, winged helix domain. (B) Interactions between ZAR1 bait constructs and 11 PBL preys were assessed in the absence and presence of HopZ1a and/or ZED1 alleles integrated at the *ho* locus of strains EGY48 (*MAT* α) or RFY206 (*MAT* **A**), corresponding to Y2H (interaction scheme A), Y3H (interaction schemes B), and Y4H (interaction scheme C) assays (see S1 Fig). The prey array layout shown at right represents PBL interactions with ZAR1^ΔCC^ in the presence of both ZED1 and HopZ1a^wt^. The colours of the labels at each array position (white or black) are determined by the relative interaction strength (see S2 Fig; Materials and Methods).

Although we have demonstrated above that ZED1 mutants with restored kinase motifs gain HopZ1a-independent binding affinity for PBL kinases, it is possible that these interactions are formed by a binding interface that differs from that required for HopZ1a-dependent PBL-ZED1-ZAR1 interactions. To address this, we integrated genes encoding ZED1^wt^ or ZED1^‘DT’^ (ZED^N173D V212T^) at the yeast *ho* locus to create a HopZ1a-independent Y3H system in which to test possible PBL-ZED1-ZAR1 interactions. Expression of ZED1^‘DT’^ allowed HopZ1a-independent interactions between ZAR1^wt^ and PBL17 / PBL9, whereas ZED1^wt^ did not (Fig 5B, compare columns 7 and 8). Moreover, in the presence of ZED1^‘DT’^ (but not ZED1^wt^), ZAR1^ΔCC^ interacts more robustly and with more PBLs (eight of the eleven tested; Fig 5B, compare columns 7 and 8 with 23 and 24), indicating that the coiled coil domain negatively regulates formation of the ZAR1-ZED1-PBL ternary complex in HopZ1a-independent contexts as well. Importantly, ZAR1^ΔLRR^ did not interact with PBL kinases in the presence of ZED1^‘DT’^ indicating that the leucine-rich repeat domain is required for both HopZ1a-dependent and HopZ1a-independent interactions.

### ZED1 mutants with HopZ1a-independent PBL-binding promote HR in *Arabidopsis*

We hypothesized that ZED1/PBL interactions promote ZAR1-mediated immunity, similar to the ZRK1/PBL2/ZAR1-mediated immunity triggered by AvrAC [12]. As such, mutant ZED1 alleles that allow HopZ1a-independent interactions with PBL kinases should also promote immunity in the absence of the effector. To test this, we transformed the *Arabidopsis* ecotype Col-0 with four dexamethasone-inducible, HA epitope-tagged ZED1 alleles (ZED1^wt^, ZED1^N173D^, ZED1^V212T^, and ZED1^‘DT’^), and examined the first generation (T1) for a dexamethasone-induced tissue collapse similar to the HR observed in plants expressing the *P. syringae* effector AvrRpt2 [37] (see column 2 in Fig 6AB, S12 Fig AB, and S13 Fig AB). We established expression profiles for at least 8 individual T1 transformants from each ZED1 allele (panel C in Fig 6, S12 Fig, S13 Fig) prior to testing for dexamethasone-inducible cell death activity. These same plants (with one leaf removed), along with untransformed Col-0 and *zar1-1* mutant [11] plants as controls, were then sprayed with dexamethasone and observed visually for signs of dexamethasone-induced HR. As expected, untransformed wild-type *Arabidopsis* (Col-0) and *zar1-1* mutant plants were both unaffected by dexamethasone treatment (columns 1 and 3 in Fig 6AB, S12 Fig AB, and S13 Fig AB). *Arabidopsis* plants expressing ZED1^V212T^ and ZED1^‘DT’^ displayed tissue collapse similar to AvrRpt2-expressing plants 72 hours after dexamethasone treatment (columns 4 and 5 in S13 Fig AB; columns 6 and 7 in Fig 6AB), whereas plants expressing ZED1^wt^ and ZED1^N173D^ did not (columns 4, 5 in Fig 6AB; columns 4-7 in S12 Fig AB). These results suggested that PBL-interacting ZED1 mutants can trigger HopZ1a-independent immunity and led us to investigate whether this response is ZAR1-dependent.

**Fig 6.**
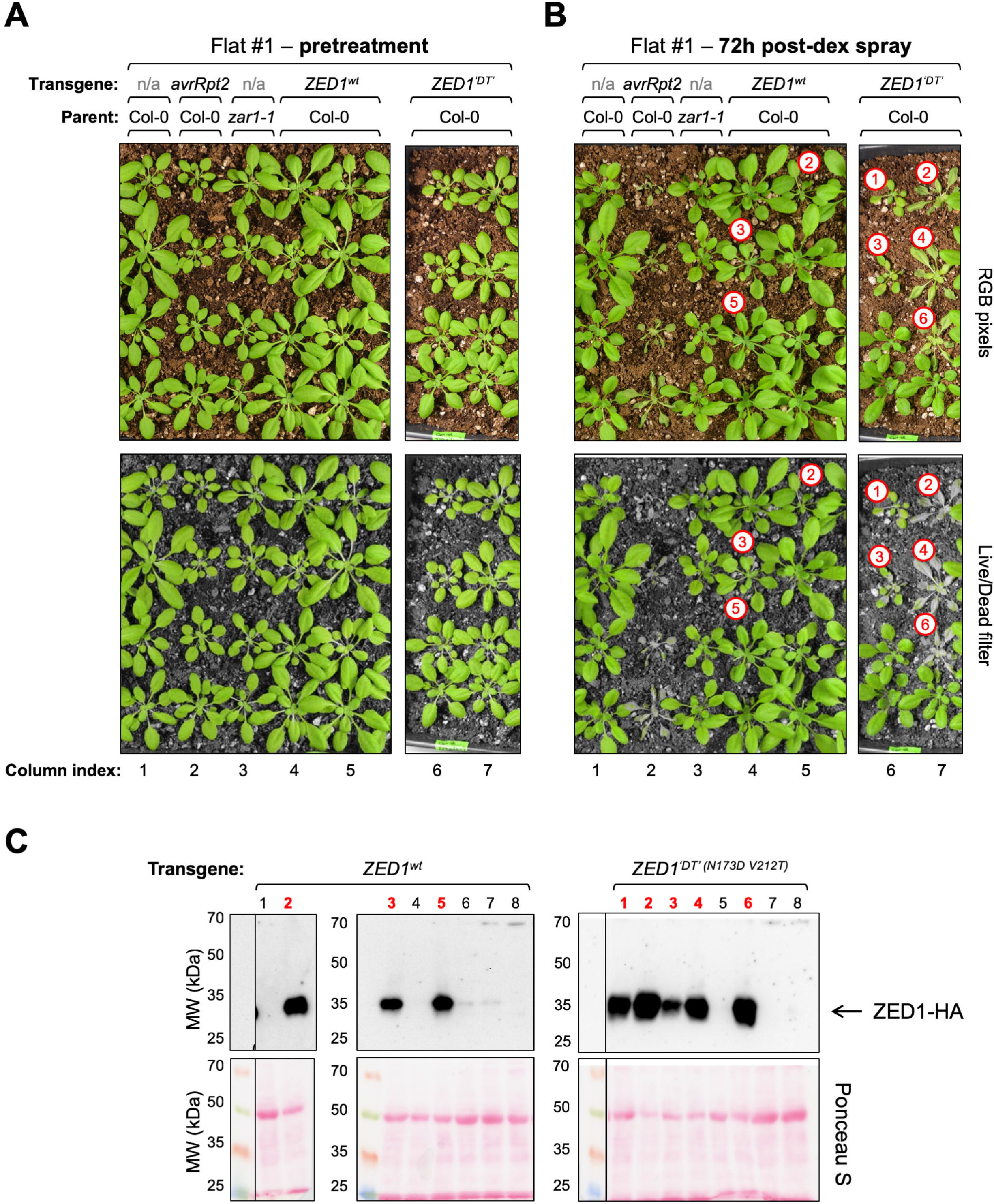
Dexamethasone-induced expression of ZED1^‘DT’^ (N173D V212T) causes whole-plant HR in *Arabidopsis*. BASTA-resistant *Arabidopsis* transformants bearing dexamethasone-inducible ZED1 alleles (columns 4-7) were tested alongside control plants (columns 1-3; untransformed Col-0, transgenic plants with dexamethasone-inducible AvrRpt2, and untransformed *zar1-1*). Photographs of control plants and *Arabidopsis* T1 transformants are shown both before (A) and ∼72 h after dexamethasone induction of transgenes (B). Panels A and B present full-colour RGB images (top) as well as a filtered representation (‘Live/Dead filter’; bottom) that converts dead/dying plant tissue and surrounding soil to grayscale pixels while leaving pixels representing healthy *Arabidopsis* tissue unchanged (see S20 Fig; Materials and Methods). Red numbered circles indicate those plants for which transgene expression was detected and correspond to the electrophoresis lane indices shown in panel C, below. Note that plants presented in this Figure were grown on the same flat and so experienced identical conditions with respect to watering, lighting, and dexamethasone sprays; whole-flat images were cropped to remove unrelated plants. (C) Parallel assessment of transgene expression in tissue from the same plants shown in panels A and B. HA-tagged transgene expression in T1 transformants is shown (top), and Ponceau S staining of total protein transferred to the nitrocellulose membrane (bottom) demonstrates consistent sample loading and protein transfer across all lanes.

### ZED1 mutants trigger HopZ1a-independent, ZAR1-dependent HR in *Nicotiana benthamiana*

*Nicotiana benthamiana* has two ZAR1 homologues (*Nb*ZAR1 and *Nb*ZAR2) that are ∼57% identical to *Arabidopsis* ZAR1 [38]. Despite lacking closely-related homologues of ZED1 and ZRKs (S14 Fig), ZAR1-dependent immune responses triggered by HopZ1a can be recapitulated in *N. benthamiana* when complemented with *Arabidopsis* ZED1 [38]. Cell death resulting from an HR is observed as early as 8 hours following induction of transgene expression in *N. benthamiana* leaves co-transformed with ZED1 and HopZ1a, but not in leaves transformed with either ZED1 or HopZ1a alone [38]. Since BLASTP searches demonstrate that *Nicotiana spp.* do have highly-similar homologues of *Arabidopsis* PBLs (S14 Fig), we hypothesized that our ZED1 mutants with HopZ1a-independent PBL binding may be sufficient to trigger ZAR1-dependent HR in *N. benthamiana*. We therefore tested the ability of ZED1 mutants described above to cause HR in *N. benthamiana* using transient transformations. As described previously [38], macroscopic HR-like cell-death was apparent by 8-24 h post-induction of protein expression in *N. benthamiana* tissue co-transformed with ZED1^wt^ and HopZ1a, but not in tissue transformed with either ZED1^wt^ or HopZ1a alone (S15 Fig). In contrast, ZED1^‘DT’^ and ZED1^V212T^ were able to cause HR in *N. benthamiana* even in the absence of HopZ1a (S15 Fig, panel A). Overexpression of wild-type ZED1 did not induce HR despite similar levels of expression as ZED1^‘DT’^ (N173D V212T; S15 Fig, panel B).

We also tested ZAR1-dependence of these effector-independent cell death phenotypes by subjecting *N. benthamiana* plants to virus-induced gene silencing (VIGS) [38–40] two weeks prior to transformation and induction of ZED1 expression. Although the HR-mediated cell death induced by ZED1^‘DT’^, ZED1^V212T^ and HopZ1a/ZED1^wt^ was unaffected in plants experiencing silencing of the negative control *GUS* gene (Fig 7A, left), the HR induced by all of these transformations was completely abolished in plants that received a *ZAR1* silencing construct (Fig 7A, right).

**Fig 7.**
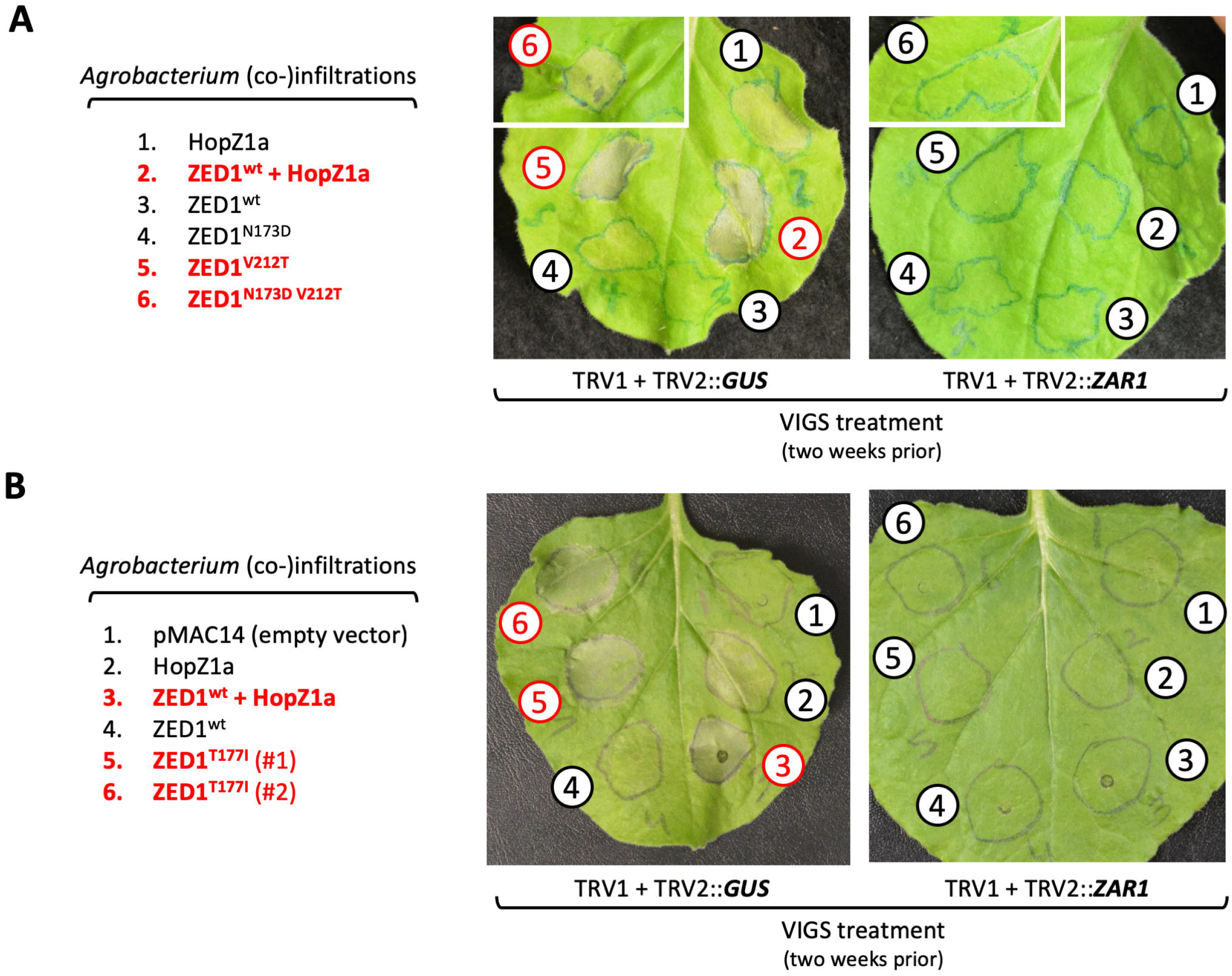
Virus-induced gene silencing of *NbZAR1* expression abolishes HopZ1a-independent HR triggered by ZED1 alleles. Leaves from *N. benthamiana* plants were infected with Tobacco Rattle Virus-derived gene silencing constructs targeting *GUS* (left) or *ZAR1* (right) two weeks prior to transformation by localized pressure-infiltration of *Agrobacterium* cell suspensions delivering the indicated dexamethasone-inducible transgenes. (A) ZED1^V212T^ substitution mutants are contrasted with ZED1^wt^ and ZED1^N173D^. Images show infiltrated leaves ∼48 h post-induction by spray with dexamethasone. Note that the images showing infiltration spot #6 are from separate leaves but from the same plants as the corresponding infiltration spots #1-5; leaf images were cropped to remove additional unrelated infiltrations. Images are representative of six different leaves infiltrated in the same way (two leaves each from three individual plants) and are consistent with multiple independent experiments. (B) A ZED1^T177I^ substitution mutant is contrasted with ZED1^wt^. Images are representative of ten different leaves infiltrated in the same way (five replicates each in two independent experiments).

To determine if pathogen-induced perturbations of ZED1 are sufficient to activate ETI, we tested whether mimicking HopZ1a acetylation of ZED1 T177 can activate plant immunity. Given that an isoleucine substitution of a PBL acetylation site acted as a gain-of-function mutant with respect to ZED1 / PBL interactions in yeast (Figure 4), we tested the ability of ZED1^T177I^ to induce HR in *N. benthamiana*. Like ZED1^V212T^, expression of ZED1^T177I^ was also capable of inducing effector-independent HR (Fig 7B; S15 Fig, panel C). Importantly, this HR was also dependent on ZAR1, supporting our hypothesis that acetylation of ZED1 by the type III effector HopZ1a activates ZAR1-mediated immunity [14].

## DISCUSSION

In this study we have examined how kinase-pseudokinase interactions induced by the *P. syringae* T3E HopZ1a contribute to the induction of plant immunity. We have shown that HopZ1a can acetylate PBL (RLCK family VII) kinases and promote their interactions with the ZED1 pseudokinase (RLCK family XII). In addition, we have also shown that ZED1 mutants that restore similarity to canonical kinase motifs gain HopZ1a-independent PBL-binding activity and can induce a ZAR1-dependent hypersensitive immune response (HR) *in planta*. Finally, we present evidence that isoleucine functions as an acetyl-threonine mimic, since mutation of a HopZ1a acetylation site on ZED1 (T177) to isoleucine also triggers ZAR1-dependent immunity. Overall our data suggest that ZED1-PBL interactions provide a sensor that can monitor for perturbations of both kinase families to trigger ZAR1-mediated ETI. The switch-like structural features of kinases - including their propensity to conditionally interact with one another - seem to have been exploited by the plant immune system to survey for effector-induced perturbations of the plant kinome.

### Functional characterization of the degenerate ZED1 active site residue, N173

One conspicuous pseudokinase feature of ZED1 - even among ZRKs - is the degenerate kinase catalytic site (N173), wherein the catalytic aspartate (acidic side chain) typically present in active kinases is replaced by asparagine (structurally-similar to aspartate, but with a basic side chain). We created the ZED1^N173D^ mutant to test the consequences of this substitution both in Y2H assays and in an inducible expression system *in planta*. Although our yeast assays revealed that ZED1^N173D^ gains effector-independent affinity for PBLs, we did not observe induction of cell death (immunity) following expression of ZED1^N173D^ in *Arabidopsis* (S12 Fig) or *N. benthamiana* (Fig 7A). However, a distinct substitution at this position (N173S; which introduces a shorter, polar side chain) was previously shown to display ZAR1-dependent developmental and immunity-related phenotypes that are conditionally observed at elevated temperature (25°C), but not at a lower temperature (18°C) [41]. Since our *in planta* experiments were carried out at ambient (room) temperatures (∼20°C), it is conceivable that some of our ZED1 mutants (such as ZED1^N173D^) may display temperature-dependent phenotypes.

### The ZED1 activation loop is implicated in PBL binding and immune activation *in planta*

Although both ZED1^N173D^ and ZED1^V212T^ mutants acquired strong HopZ1a-independent PBL binding activity, only the ZED1^V212T^ and ZED1^N173D V212T^ mutants (with restored activation loops) were sufficient to induce immunity *in planta* (Fig 6; S13 Fig; Fig 7A; S15 Fig). ZED1 is unique among the ZRKs in that it lacks any candidate phospho-accepting residues in the activation loop (S6 Fig, panel B). This feature is not specific to the Col-0 ecotype of *Arabidopsis*, since examination of 813 unique ZED1 sequences representing more than 1000 distinct and globally-distributed ecotypes identified only four with potential phospho/acetyl-accepting sites between the imperfect ‘DFG’ and ‘APE’ motifs that define the activation loop (S16 Fig; see Materials and Methods).

### Activation loop-mediated interface between ZRKs and PBLs

The HopZ1a acetylation target S244 of PBS1 (Fig 1B) provides further evidence for the functional importance activation loops in kinase-pseudokinase interactions. Glutamine substitution of this residue blocks HopZ1a-induced ZED1 binding activity, whereas an isoleucine substitution of the homologous activation loop position in PBL15 (S260I) acts as a gain-of-function mutant that mimics HopZ1a-induced ZED1-PBL binding (Fig 4B). These results suggest that kinase-pseudokinase (PBL-ZRK) interactions can be promoted by activation loop-dependent structural changes in either of these protein families.

We therefore speculated that an interface between ZED1 and PBLs involving the sites implicated by acetylation and/or mutational analyses may be similar to the previously described structure of a mutant IRAK4 kinase domain [42]. This mutant (bearing a catalytic site aspartate to asparagine substitution) gained the ability to dimerize in solution and was crystalized as an asymmetric dimer mediated by interactions between activation loops [42]. Remarkably, homology modeling that superimposes ZED1 and PBS1 on the two monomers of the asymmetric IRAK4 homodimer indicates that such a ‘front-to-front’ interface positions the acetylated PBL activation loop residue (i.e. PBS1 S244 or PBL15 S260) in very close proximity to ZED1 residues implicated by our mutational analysis (N173, T177, and V212; S17 Fig). This hypothetical ZED1-PBL interface is corroborated by recent cryo-EM-derived structures of ZAR1-ZRK1-PBL2 complexes; both a nucleotide-free intermediate form [17] and the pentameric, ATP-bound, activated resistosome [18] feature pseudokinase-kinase interfaces where the uridylated PBL2 activation loop residues are closely associated with the ZRK1 activation loop and occupy the cleft between the amino-terminal and carboxy-terminal pseudokinase subdomains (S18 Fig, S19 Fig).

Kinase activation loops are also targeted by other T3Es. Cleavage of PBS1 by the T3E HopAR1 occurs after the lysine (K243) immediately preceding S244 [8], a modification that is perceived by the NLR RPS5 [9]. AvrAC uridylates serine and threonine residues in the activation loops of PBL2, BIK1 and RIPK [43,44], similar to the acetylation of PBS1 by HopZ1a. These conserved AvrAC uridylation sites are required for PBL2 interactions with ZRK1/RKS1 [12], and are just two positions ‘downstream’ of PBS1 S244 (S6 Fig, panel B). In human cells, YopJ (a HopZ-related effector from the human plague pathogen, *Yersinia pestis*) acetylates the activation loops of MAP kinase kinases (MEK2, MAPKK6) and both the α and β subunits of the IκB kinase (IKK) complex, resulting in inhibition of MAP kinase and NF-κB signaling pathways [45,46]. It has long been established that the phosphorylation status of activation loops has a role in influencing conformational changes that both position catalytic components and regulate access of substrates to the catalytic site [47,48]. As such, kinase activation loops may represent attractive targets for invading pathogens, while also providing the structural influence required to sense and transduce kinase domain perturbations.

### ZAR1 recruitment

Our yeast four-hybrid data indicate that ZAR1 forms a ternary complex with ZED1 and PBL kinases that is stimulated by the presence of HopZ1a. Formation of this ternary complex is dependent on ZED1 (Fig 5) and is likely similar to the ZAR1-ZRK1-PBL2 complexes formed as a result of uridylation of PBL2 by AvrAC [18]. Notably, an intact ZAR1 LRR domain is required for both ZAR1-ZRK1 interactions [12] and also for formation of the ZAR1-ZED1-PBL ternary complex promoted by HopZ1a (Fig 5), consistent with co-immunoprecipitation experiments in *N. benthamiana* with *Arabidopsis* ZED1 and ZAR1 truncation mutants [38] and with the ZAR1-ZRK1 interface observed in cryo-EM structures [17,18]. These findings are distinguished from the PBS1/RPS5 model wherein PBS1 interacts with the coiled-coil (CC) domain of RPS5 until cleavage of the PBS1 activation loop by HopAR1 abolishes this interaction, suggesting that in the case of RPS5, binding by PBS1 is required to restrain an inherently active NLR [9]. Although direct interactions between ZED1 and the ZAR1 CC domain have been previously reported [14,38], our data suggest indicate that the CC domain of ZAR1 can negatively regulate formation of a ZAR1-ZED1-PBL ternary complex (Fig 5B). Indeed, inactive ADP-bound ZAR1-ZRK1 complexes are stabilized by CC-mediated interdomain contacts with the HD1, WHD, and LRR domains [17] - dramatic repositioning of the NBD and HD1 domains and a fold-switch in the CC domain are required for ADP/ATP exchange and subsequent assembly of the resistosome [17,18].

Overall, our results support models suggesting that acetylation of ZED1 and/or of PBL kinases by HopZ1a promotes formation of similar ternary complexes with ZAR1; this process requires the LRR domain and is likely to be negatively-regulated by the ZAR1 CC domain (Fig 8, models B, C). Ternary complex formation may occur by stabilizing a preformed ZAR1-ZED1 complex, as described for the ZAR1-ZRK1-PBL2 interactions [12,17,18]. Alternatively, HopZ1a-stabilized ZED1/PBL dimers may bind ZAR1 as a pre-formed unit to activate immunity, since we have found that HopZ1a-induced ZED1-PBL interactions are ZAR1-independent (Fig 2A; Fig 4B; S8 Fig; S9 Fig; S10 Fig).

**Fig 8.**
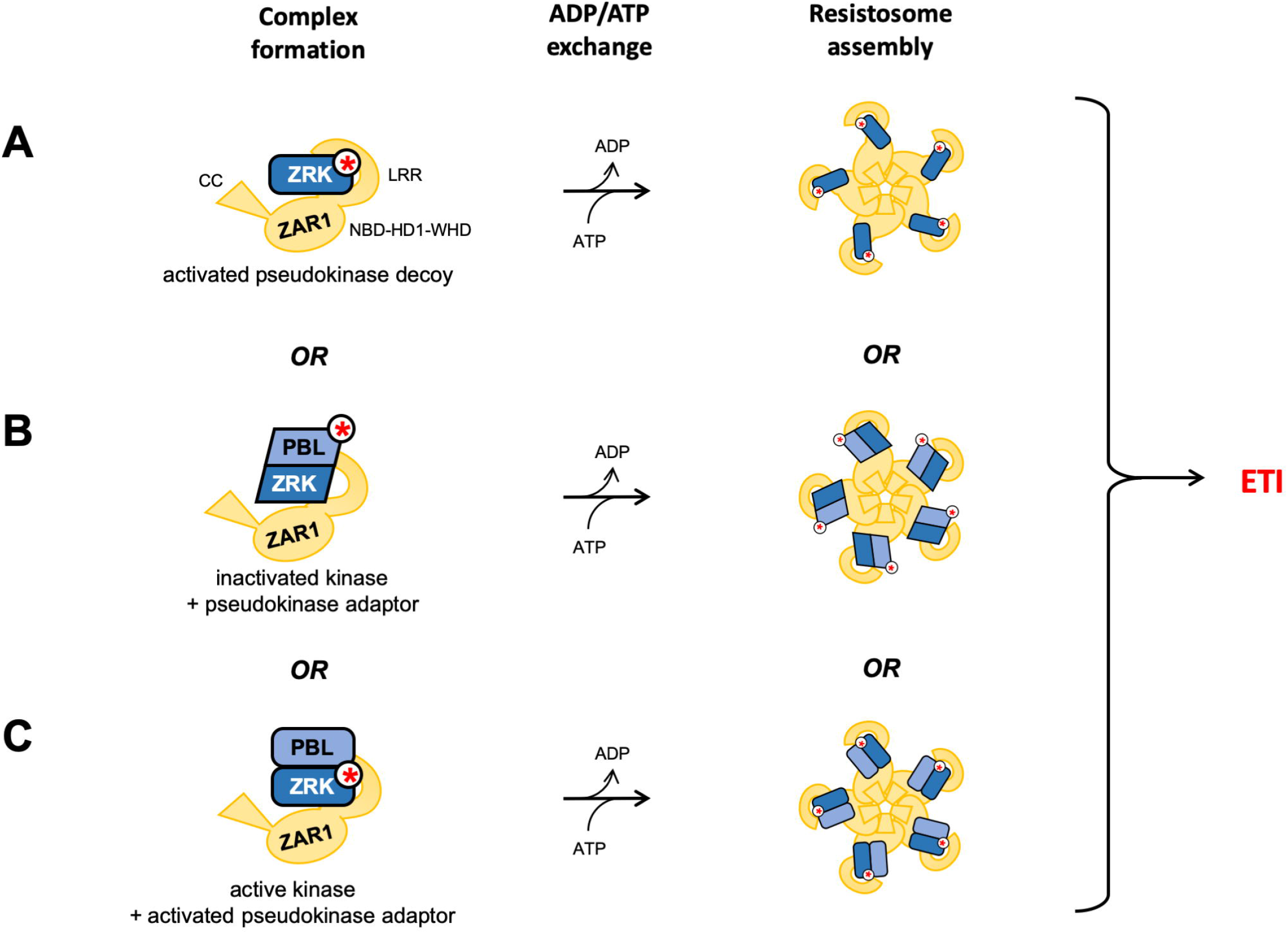
Alternate models for explaining activation of ZAR1-dependent immunity in *Arabidopsis*. ZAR1 interacts with pseudokinases and kinases to achieve ADP/ATP nucleotide exchange and assembly of a resistosome. Active or ‘active-like’ kinase domain conformations are represented with rounded rectangles, while inactive and ‘inactive-like’ kinase domain conformations are represented with parallelograms. (A) In the decoy model, post-translationally-modified ZRKs are presumed to adopt an ‘active-like’ conformation that is required to activate ZAR1 and trigger ETI. (B) In the adaptor model proposed by Wang et al. [12], post-translational modification of PBLs results in conversion to an inactive conformation that is sensed by ZRKs to promote nucleotide exchange by ZAR1 and activation of ETI. (C) An alternative adaptor model suggested by our data implies that post-translational-modification of ZRK pseudokinases may be sufficient to recruit PBLs for ZAR1 activation by inducing an ‘active-like’ pseudokinase conformation. In other words, models B and C both allow heterodimer formation only when kinase and pseudokinase have adopted similar structural conformations – inactivated kinases bind to ‘inactive-like’ pseudokinases, but similarly, modified pseudokinases adopting ‘active-like’ conformations as a result of post-translational modifications can also bind to active (unmodified) PBLs to activate ZAR1.

### Models for ZAR1 activation by HopZ1a

According to our original decoy model of ZED1 function, acetylation of ZED1 should be sufficient to activate ZAR1 and trigger immunity [14] (Fig 8, model A). In this report however, we have described HopZ1a-induced ZED1-PBL (and ZAR1-ZED1-PBL) interactions that are consistent with the adaptor model proposed by Wang *et al.* [12] and recent structures of a ZAR1-ZRK1-PBL2^UMP^ resistosome [17,18] (Fig 8, model B), suggesting that the ability to conditionally interact with PBLs may be a general feature of ZRK pseudokinases. Nevertheless, our experiments have not ruled out a role for effector-mediated modification of ZED1 in the activation of ZAR1. Indeed, HopZ1a can acetylate ZED1 at sites proximal to kinase motifs of known functional importance (S7 Fig, panel A), glutamine substitution of the ZED1 acetylation site T177 abolishes HopZ1a-dependent ZED1-PBL binding (S10 Fig, panel B), and isoleucine substitution of T177 is sufficient to cause ZAR1-dependent induction of immunity in *N. benthamiana* (Fig 7B). We speculate that, like ZRK1 [17,18], ZED1 may act as a nucleotide exchange factor for ZAR1 to activate immunity, although in this case nucleotide exchange-promoting activity may be activated by direct acetylation of ZED1. T177 is located between the degenerate catalytic site residue (N173) and a trio of residues contributing to the ‘catalytic spine’ (I179, F180, I181), and its modification by HopZ1a may influence kinase structure by repositioning catalytic and regulatory spines and/or the activation loop (S19 Fig) [21]. Such structural rearrangements may allosterically regulate the relative orientations of ZAR1 subdomains to promote ADP/ATP exchange. PBL interactions may also be required for ZED1-dependent activation of ZAR1, since HopZ1a activity can promote ZED1-PBL binding (Fig 2A) and formation of ZAR1-ZED1-PBL ternary complexes (Fig 5). Binding of PBL kinases to (acetylated) ZED1 might further stabilize conformational changes required to induce ADP/ATP exchange by ZAR1, resistosome assembly, and immune activation.

Although PBLs are also acetylation targets of HopZ1a (Fig 1), ZED1 mutations that restore degenerate pseudokinase motifs can also promote HopZ1a-independent ZED1-PBL interactions and ZAR1-ZED1-PBL complex formation (Fig 3B; S8 Fig; S9 Fig, Fig 5B). We speculate that these mutations allow ZED1 to adopt an activated pseudokinase structure similar to that of acetylated ZED1. We therefore introduce an additional model for activation of ZAR1, recognizing that effector-mediated modification of either ZRKs or PBL kinases may be sufficient to form kinase-pseudokinase dimers and activate ETI (Fig 8, models B and C). We hypothesize that T3E-induced modifications introduce structural changes in ZRKs that can recruit unmodified PBL kinases to activate ZAR1 (Fig. 8, model C). Together with the ZRK1/PBL2 adaptor model (Fig. 8, model B), this new model suggests that weak basal kinase-pseudokinase interactions are enhanced by structural transitions of either binding partner between complementary active/inactive conformations. The recently-described ZAR1-ZRK1-PBL2 complexes [18] may indeed reflect a structural inactivation induced by post-translational modification (PTM), since the structure of PBL2 is only partly defined, and its B-factors are elevated compared to those for ZRK1 (S18 Fig) and ZAR1. In contrast, while the ‘catalytic’ and ‘regulatory’ spines of ZRK1 do not appear to undergo conspicuous structural changes in response to PBL2 binding or ZAR1 nucleotide occupancy, binding of PBL2^UMP^ stabilizes the activation loop of ZRK1 (S19 Fig) [17,18].

The PTMs catalyzed by HopZ1a (acetylation; adds 42 Da) and AvrAC (uridylation; adds 324 Da) differ in size and in physicochemical properties, and the diversity of enzymatic functions possessed by these and other effectors likely represents a significant force contributing to the evolutionary pressures driving diversification of the ZRKs. We note that by using pseudokinase adaptors that monitor kinase domain conformations rather than specific PTMs plants would be able to recognize an even greater diversity of potential microbial effectors with diverse enzymatic activities. Further structural characterization of ZRK pseudokinases (both alone and in complex with PBLs and/or ZAR1) will be important for critical assessment of our model presenting ZRK/PBL modules as molecular switches that are sensitive to perturbations of kinase structure.

### Functional significance of the PBL kinases targeted by HopZ1a

It is likely that the PBL kinases with HopZ1a-dependent ZED1 binding include virulence targets that are manipulated to promote *P. syringae* pathogenesis. Notably, a number of PBLs have been implicated in plant immunity, including BIK1 [49–52], RIPK [53,54], PBL13 [55], and PBL27 [56,57], and we have shown that the latter two participate in HopZ1a-dependent interactions with ZED1 (Fig 2A). PBL kinases were not identified in our earlier forward genetic screen for loss of HopZ1a-induced HR [14] (in contrast to the susceptibility of *Arabidopsis pbl2* mutants to infections with AvrAC-expressing *X. campestris* [12]). If PBLs do in fact play a role in activation of ZAR1-mediated ETI by binding to (acetylated) ZED1, we expect that there are multiple PBLs capable of fulfilling this function. Functionally-redundant PBL kinases would as result be individually dispensable for HopZ1a-induced ETI, or alternatively, one or more of these PBLs may also be essential for viability, precluding their recovery as loss-of-function mutants.

### Functional diversity of HopZ alleles

HopZ1a is a member of a large and diverse family of bacterial effector proteins with similarity to the YopJ effector from *Y. pestis*. YopJ-like effectors are produced by a variety of pathogens of both plants and animals, and even among *P. syringae* strains there are five recognized distinct lineages of HopZ effectors (HopZ1 through HopZ5) [58,59]. This inter-allelic sequence variation confers significant functional differences, since only HopZ1a is capable of inducing ETI in *Arabidopsis*. HopZ1b, the allele most closely-related to HopZ1a (64% identity), can trigger a weak, HR-like tissue collapse, but this response is ZAR1-independent and is only observed in ∼25% of infiltrated leaves [11,60]. HopZ1b induced only a subset of the PBL-ZED1 interactions promoted by HopZ1a, consistent with its inability to activate robust ZAR1-dependent immunity. The PBL kinases that interact with ZED1 specifically (or more strongly) in the presence of HopZ1a than HopZ1b thus represent promising candidates for key regulators of ZAR1 activation by HopZ1a. HopZ2 (26% identical to HopZ1a) did not promote interactions between ZED1 and PBL kinases (S5 Fig) and is able to promote *P. syringae* virulence in *Arabidopsis* without activating ZAR1 immunity [60]. Likewise, HopZ3 (23% identical to HopZ1a) does not activate ZAR1, however it is able to interact with PBLs (RIPK, PBS1, BIK1, and PBL1) and can also acetylate the activation loop of RIPK (S251 and S252 - identical to the sites uridylated by AvrAC) to dampen ETI mediated by the NLR RPM1 [61]. In our assays, however, co-expression with HopZ3 did not confer ZED1 binding activity upon RIPK or any other PBL (S5 Fig). Overall, the promotion of most ZED1-PBL interactions is specific to HopZ1a/HopZ1b, suggesting that other HopZ alleles may have adopted distinct modification strategies to avoid recognition of their modified substrates by ZED1 and ZAR1 in *Arabidopsis*.

Interestingly, a related HopZ-like T3E from *Xanthomonas perforans*, XopJ4/AvrXv4 (25.5% identical to HopZ1a across 337 non-gapped sites) triggers a ZAR1-dependent ETI in *N. benthamiana* that is dependent on an RLCK XII protein named XOP<u>J</u>4 <u>IM</u>MUNITY <u>2</u> (JIM2) [62]. ZED1 is the *Arabidopsis* protein most closely-related to JIM2 (BLASTP E-value of 5e-73; 41% identity spanning 92% of the JIM2 query sequence) and notably, JIM2 has degenerate kinase motifs (including a ‘dead’ HRD catalytic motif - ‘YRI’), suggesting that *Nb*ZAR1 may also use a pseudokinase/kinase module to detect perturbations of *N. benthamiana* signaling [62].

### Concluding Statement

Kinases and pseudokinases possess switch-like structural features that are often allosterically regulated by post-translational modifications (PTMs) to influence their activity (in the case of active kinases) and/or alter their protein interaction profiles, resulting in reorganization and redistribution of signaling network activities. The plant immune system appears to have harnessed these switch-like features to detect kinase/pseudokinase perturbations that are induced by pathogen-delivered effector proteins. Modifications of either the ZED1/ZRK pseudokinases (RLCK XII) or of PBL kinases (RLCK VII) can promote interactions between these two families, as well as subsequent/concomitant interactions with the NLR ZAR1 to activate ETI. ZRK/PBL interactions are likely determined by a structural switch that is flipped by effector-mediated PTMs (Fig 8), providing an array of discriminating sensors that can detect diverse perturbations of the plant kinome (including both kinases and pseudokinases). Further investigations of these intermolecular interactions, the ways in which they are influenced by bacterial effectors, and the structural features required of these components for robust immunity signaling will together provide valuable insights into the molecular mechanisms underlying pathogen perception by plant immune systems.

## MATERIALS AND METHODS

### Construction of plasmids for yeast two-hybrid assays

PBL coding sequences were obtained from the ABRC (where available) as pENTR223 or pUNI51 clones (Supplementary Table 1). Additional PBL coding sequences were synthesized by General Biosystems, Inc. (USA) as Gateway™-compatible pDONR207 clones. Recombination reactions using ‘LR clonase’ (Invitrogen) were used to shuttle coding sequences into a Gateway-compatible derivative of the prey plasmid pJG4-5 (which encodes HA-tagged amino-terminal fusions of the B42 activation domain with nuclear localization sequences; NLS-B42-HA-prey).

Donor plasmids carrying truncations of ZAR1 lacking the LRR (ΔLRR, nucleotides 1-Δ 1545, amino acids 1-515) or coiled-coil (ΔCC, nucleotides 433-2556, amino acids 145-852) Δ domains, or carrying isolated individual domains (CC, nucleotides 1-432, amino acids 1-144; NB, nucleotides 433-1545, amino acids 145-515; LRR, nucleotides 1546-2556, amino acids 516-852) were prepared by PCR amplification and recombination into pDONR207 with BP clonase (Invitrogen). Subsequent LR reactions were used to shuttle these constructs into the bait plasmid pEG202 (which encodes an amino-terminal LexA DNA-binding domain; LexA-bait). Oligonucleotide primers used for site-directed mutagenesis are indicate in S5 Table. Plasmids were introduced into yeast strains according to standard LiAc/PEG methods [63,64] following sequence verification by sequencing both DNA strands of the entire coding sequences and across cloning junctions for each clone of interest, using Sanger sequencing services provided by the Center for the Analysis of Genome Evolution and Function (CAGEF, University of Toronto).

### Co-expression/immunoprecipitation of PBS1 and ZED1 with HopZ1a in yeast and sample preparation

*PBS1* was shuttled from pDONR207 into the centromere-based yeast plasmid pBA350V [65] using LR clonase to create a galactose-dependent expression vector for production of PBS1-FLAG in yeast. pBA350V::*PBS1-FLAG* was introduced into a derivative of yeast strain Y7092 with chromosomally-integrated *hopZ1a-FLAG* at the *ho* locus, as described previously [14,66].

FLAG-tagged proteins were expressed in yeast, and cell lysates were prepared and probed with anti-FLAG-conjugated agarose resin (Sigma) as described previously [14,66]. FLAG-tagged proteins immunoprecipitated in this manner were eluted by incubating with 100 uL of FLAG peptide solution (150 ug/mL FLAG peptide in TBS) for one hour at 4°C. Eluted material was then dried to a pellet under vacuum and stored at −80°C prior to subsequent mass spectrometry analysis. Dried protein samples were re-solubilized in 50 mM ammonium bicarbonate (pH 7.8) and then subjected to reduction with dithiothreitol at 56°C, alkylation with iodoacetamide at room temperature, and overnight digestion with sequencing-grade trypsin (Promega) at 37°C. This enzymatic reaction was terminated by addition of formic acid (to 3%), and digestion products were purified and concentrated with Pierce C18 spin columns (Thermo Fisher Scientific), then again dried to a pellet under vacuum. Peptide samples were then solubilized in 0.1% formic acid prior to LC-MS/MS analyses.

### LC-MS/MS analysis of immunoprecipitated proteins - chromatography and mass spectrometry

Subsequent analytical separation was performed on a homemade, 75 μm i.d. column (New Objective, Woburn, MA) gravity-packed with 10 cm of 100 Å, 5 μm Magic C18AQ particles (Michrom, Auburn, CA). Peptide samples were loaded onto the analytical column using a variable gradient with a flow rate of 300 nL/min. The gradient utilized two mobile phase solutions: A - water/0.1% formic acid; and B - 80% acetonitrile/0.1% formic acid. Samples were analyzed on a linear ion trap-Orbitrap hybrid analyzer outfitted with a nano-spray source and EASY-nLC 1200 nano-LC system. The instrument method consisted of one MS full scan (400– 1400 *m/z*) in the Orbitrap mass analyzer, an automatic gain control target of 500,000 with a maximum ion injection of 500 ms, one microscan, and a resolution of 60,000. Six data-dependent MS/MS scans were performed in the linear ion trap using the three most intense ions at 35% normalized collision energy. The MS and MS/MS scans were obtained in parallel fashion. In MS/MS mode automatic gain control targets were 10,000 with a maximum ion injection time of 100 ms. A minimum ion intensity of 1000 was required to trigger an MS/MS spectrum. The dynamic exclusion was applied using an exclusion duration of 145 s. Each sample was analyzed in triplicate. A spectral library was created using Proteome Discoverer 2.0 (Thermo Fisher Scientific).

### Protein identification and database searches

Proteins were identified by searching all MS/MS spectra against a large database composed of the complete proteome of *Saccharomyces cerevisiae* strain S288C (ATCC 204508; UniProt proteome ID UP000002311) supplemented with sequences for *P. syringae* HopZ1a (WP_011152901.1), and the *Arabidopsis* kinases PBS1 (NP_196820.1) or ZED1 (NP_567053.1) using SEQUEST [67]. A fragment ion mass tolerance of 0.8 Da and a parent ion tolerance of 30 ppm were used. Up to two missed tryptic cleavages were allowed. Methionine oxidation (+15.99492 Da), cysteine carbamidomethylation (+57.021465 Da), and acetylation (+42.01057 Da; for serines and threonines only) were set as variable modifications. Additional description of these data is provided in S1 File.

### Construction of yeast strains enabling three-hybrid and four-hybrid interaction analyses

Derivatives of plasmid pBA2262 used to generate yeast strains with chromosomally-integrated copies of *hopZ1a^wt^, hopZ1a^C216A^, hopZ1b, hopZ2* and *hopZ3* have been described previously [66]. Briefly, genes cloned into pBA2262 are under the control of the *GAL* promoter (which is positively regulated by galactose), they are linked to the downstream *NAT^R^* gene (which provides resistance to nourseothricin, also known as clonNAT), and are flanked by 5’ and 3’ fragments of the *HO* gene (which encodes the homing endonuclease required for mating-type switching in wild yeast but is dispensable and inactivated - i.e. *hoΔ* - in domesticated laboratory strains). pBA2262 cannot replicate in yeast so pBA2262 derivatives are linearized by digestion with the restriction enzyme NotI prior to transformation, and chromosomal integrants resulting from double recombination events that replace the endogenous *hoΔ* locus with the gene of interest are isolated by selection on YPDA plates (yeast extract, peptone, dextrose, adenine sulfate) containing clonNAT at 100 µg/mL.

The *avrAC* coding sequence was amplified from pUC19-35S-*avrAC*-HA (plasmid generously provided by Jian-Min Zhou; Chinese Academy of Sciences, Beijing) to first create pDONR207-*avrAC* with BP clonase (Invitrogen), which in turn allowed subsequent creation of pBA2262-*avrAC* with LR clonase (Invitrogen). Similarly, *ZED1^wt^* and *ZED1^N173D V212T^* were shuttled from pDONR207-based plasmids to pBA2262 using LR clonase.

### Assessing bait-prey interactions in yeast

The haploid yeast strain EGY48 (‘alpha’ mating type; i.e. *MAT* α) and derivative strains bearing chromosomally-integrated additional genes (encoding bacterial effectors or alleles of the *Arabidopsis* pseudokinase ZED1) were transformed with query bait plasmids (pEG202 derivatives; *HIS^+^*), and transformants were isolated by selecting for prototrophy on Synthetic Defined (SD) minimal media containing 2% glucose and lacking histidine (SD +Glc -His). Similarly, haploid yeast strain RFY206 (‘A’ mating type; i.e. *MAT* **A**) carrying the *lacZ*-bearing reporter plasmid pSH18-34 (*URA*^+^) and a derivative strain bearing chromosomally-integrated *ZED1^wt^* were both transformed with prey plasmids (pJG4-5; *TRP*^+^), and transformants were isolated by selecting for prototrophy on SD glucose media lacking uracil and tryptophan (SD +Glc -Ura -Trp). 12 x 8 arrays of bait (*MAT* α) and prey (*MAT* **A**) transformants were first arranged manually on appropriate selective media, but all subsequent array manipulations were performed using a 96-pin replicator. Diploid strains carrying bait, prey and reporter plasmids were created by co-incubation of bait and prey arrays on YPDA media (8-18 h), followed by two selections on SD glucose media lacking histidine, uracil and tryptophan (SD +Glc -His -Ura -Trp). Bait-prey interactions were assessed on SD minimal media reporter plates containing 2 % raffinose, 1 % galactose, sodium phosphate (0.05 *M*, pH=7.0), X-gal (10 mg/mL) and lacking histidine, uracil and tryptophan (SD +Raf +Gal +X-gal -His -Ura -Trp). All yeast plates contained 2 % agar and were incubated at 30°C.

### Acquisition of yeast plate images and image processing

Reporter plates were imaged from the bottom using a flat-bed scanner (Epson) against a black felt background for contrast. The graphical summaries of the yeast interaction data shown in Fig 1-3, Fig 5, S4 Fig, S5 Fig, S8 Fig, S9 Fig, and S11 Fig were prepared by finding the average colour for an 81-pixel square (i.e. 9 pixels x 9 pixels) at the center of each colony. Averaged pixel intensities from yeast interaction plates were extracted by defining array positions with an interactive interface implemented in PYTHON (www.python.org) using PROCESSING (www.processing.org; py.processing.org). Colours of text labels (white or black) for the prey array interaction summaries accompanying yeast reporter plate images (and for the leaves on the phylogenetic tree shown in Fig 3B) were determined according to a threshold based on the relative intensities of the red, green and blue channels for the 9 x 9 averaged pixel: specifically, the value of the ratio blue/(red + green) was evaluated at each PBL array position, and positions where this ratio was ≥ 0.727 (relative interaction strength ≥ 0.455 i.e. mostly blue; strong interactions) were assigned white labels, while all other positions were labeled in black (S2 Fig). This value was used as a working threshold for distinguishing strong interactions from the background since it bisects the path described by the total interaction dataset presented in this study (S2 Fig). Pixel plotting and analysis was implemented in PYTHON using MATPLOTLIB [68]. Relevant scripts are available at https://github.com/DSGlab/Yeast-Array-Analysis.

### Creation of plant expression vectors

For construction of dexamethasone-inducible plant expression vectors, we cloned coding sequences (lacking stop codons) for *hopZ1a* and for wild-type and mutant *ZED1* alleles into pMAC14, which includes a dexamethasone-responsive promoter upstream of a multiple cloning site, as well as linked genes encoding resistance to kanamycin and glufosinate (also known as BASTA) [69]. *hopZ1a* and *ZED1* alleles were cloned between XhoI and StuI restriction sites present in the pMAC14 multiple cloning site, placeing the coding sequence in frame with a downstream, HA-tag-encoding sequence (and stop codon). *Agrobacterium* strain GV3101 was transformed according to standard methods [70], and transformants were selected by screening for plasmid-encoded kanamycin resistance.

### Plant transformations and expression assays

We used the floral dip method [71] to generate germ-line transformants of *Arabidopsis* using *Agrobacterium* strains transformed with pMAC14 derivatives. *Arabidopsis* transformants were selected by germinating seeds on soil infused with 0.1% BASTA (v/v in water). Transformed (BASTA-resistant) seedlings were then transplanted into fresh soil free of herbicide along with control wild-type *Arabidopsis* (Col-0), third-generation (T3) AvrRpt2-expressing plants, or *zar1-1* mutants. Transgene expression was induced by spraying each flat with ∼ 30 mL of 20 μM dexamethasone in ddH_2_O with 0.01% surfactant Silwet-L77.

Leaves of *N. benthamiana* plants were locally transformed by pressure infiltration with suspensions of pMAC14-transformed *Agrobacterium* strains using a needleless syringe applied to the underside of the leaf. Transgene expression was induced by spraying whole leaves with 20 μM dexamethasone in water with 0.01% surfactant Silwet-L77 (8 - 24h following infiltrations). Gene silencing was performed as described previously [38–40], using *Agrobacterium* strains generously provided by Maël Baudin and Jennifer D. Lewis (University of California, Berkeley).

### Preparation of protein extracts from plant tissue and immunoblot analysis

Expression of transgenes in *Arabidopsis* was assessed by punching three leaf cores from a single leaf which were then floated on 20 μM dexamethasone solution (in ddH_2_0) for ∼18h. Expression of transgenes in *N. benthamiana* was induced by spraying infiltrated plants with a solution of 20 μM dexamethasone in ddH_2_0 with 0.01% Silwet L-77. For each transformation of interest, tissue was harvested by punching three circular leaf cores 5 mm in diameter. These tissue samples were then flash-frozen in liquid nitrogen before manual grinding with a mini-pestle in a 1.5 mL Eppendorf tube. Powdered frozen plant tissue was then suspended in 100 µL of Grinding Buffer (40 mM Tris pH=7.5, 150 mM NaCl, 1 mM EDTA, 5 mM DTT, 1% Triton X-100, 0.1% sodium dodecyl sulfate) supplemented with a 1:500 dilution of plant protease inhibitor cocktail P9599 (Sigma). 15 uL from each of these extracts was then resolved by electrophoresis through 10 % polyacrylamide SDS-PAGE gels prior to protein transfer to nitrocellulose membranes. Membranes were blocked with 5% powdered skim milk solution in TBST (50 mM Tris-Cl, pH 7.5; 150 mM NaCl; 0.05% Tween-20). Primary (rat anti-HA, mouse anti-LexA, and mouse anti-FLAG) and secondary antibodies (alkaline peroxidase-conjugated goat anti-mouse) were both diluted 1:10,000 in TBST with 3% powdered milk.

### Phylogenetic analysis of kinase domains

Sequences for 46 PBL kinases and 12 ZED1-related pseudokinases were retrieved from TAIR (www.arabidopsis.org), aligned with MUSCLE [72], and trimmed to remove variable-length amino-terminal and carboxy-terminal sequences flanking the conserved kinase domain. These isolated kinase domain sequences were then supplemented with sequences from structurally-characterized kinase domains of the *Arabidopsis* receptor kinases BRI1 (PDB: 5LPV) [33] and BAK1 (PDB: 3UIM) [34], the tomato Pto kinase (PDB: 3HGK) [73], and the human interleukin-1 receptor-associated kinase IRAK4 (PDB: 4U97) [42]. These 62 sequences were then realigned with MUSCLE and any amino-terminal or carboxy-terminal sequences flanking the kinase domain were again manually trimmed. The resulting alignment is provided as a supplementary file (S2 File) and was used as input for phylogenetic inference using PhyML [74,75]. Confidence in nodes of the resulting phylogenetic tree was assessed using the approximate Likelihood Ratio Test (aLRT) [76,77]. A representation of the resulting tree (rooted on IRAK4 and including only those nodes with greater than 70% confidence) is presented in S6 Fig, panel A, and the input tree is provided as a supplementary file (S3 File). A derivative of the same tree is also shown in Fig 3B and is the result of a pruning step to remove any non-PBL leaves. Kinase domain trees were visualized using iTOL [78].

### Analysis of ZED1 activation loop sequences from diverse *Arabidopsis* ecotypes

The *ZED1* sequences for available *Arabidopsis* ecotypes were derived from DNA SNP information curated by the 1001 genomes project (http://1001genomes.org/) [79] and were annotated using MAKER (http://www.yandell-lab.org/software/maker.html) [80]. 1121 translated ZED1 sequences were collapsed into 813 unique sequences and aligned using MUSCLE [72] (S4 File). 25 of these unique sequences represent more than one *Arabidopsis* ecotype, while the remaining 788 unique sequences are ‘singletons’ representing a single *Arabidopsis* ecotype (S5 File). Although confidence in translated ZED1 sequences depends on DNA sequence quality (which varies among individual sequenced ecotypes), 92.5% of the unique ZED1 sequences (mean un-gapped sequence length of 332.7 amino acids) had fewer than 38 positions translated as an ‘X’ due to ambiguous DNA sequence information (i.e. ≤ 11.4% of the mean sequence length), and 76.3% of unique sequences had fewer than 23 positions translated as X (≤ 6.9% of the mean sequence length; S16 Fig, panels A, B, top left). A cropped alignment consisting of only the activation loop sequences (mean un-gapped sequence length of 29.0 amino acids) had a similar distribution of X position prevalence, with 89.0% of activation loop sequences containing two or fewer X positions (≤ 6.9% of mean activation loop length; S16 Fig, panels A, B, bottom left). In contrast, the distributions of serine and threonine positions (S+T) were markedly different when comparing the activation loop region with full-length protein sequences. While 88.6% of the unique *Arabidopsis* ecotype ZED1 sequences have at least 36 serine or threonine positions (≥ 10.8% of the mean sequence length) across their full length (S16 Fig, panels A, B, top right), the vast majority of ecotypes (99.5%) lack any serine or threonine residues when considering activation loop sequences alone (S16 Fig, panels A, B, bottom right). Notably, the four ecotypes with threonines in their activation loop sequences are all outliers with respect to sequence quality, with either three or six ambiguous positions translated as X (sequence ambiguity that is representative of ≤ 7% of all unique sequences; S16 Fig), so it is possible that these rare apparent phospho/acetyl-accepting activations residues are in fact artefacts resulting from poor sequence quality.

### Molecular modeling

All three-dimensional protein structural models were prepared using the interface to MODELLER [81] provided within UCSF CHIMERA [82]. Model templates were selected based on high-scoring BLASTP hits in the PDB database. The ZED1 model presented in S7 Fig and S10 Fig was constructed using the multiple template option of MODELLER and is based on homology with both *Solanum pimpinellifolium* (tomato) Pto kinase (PDB: 3HGK; 30% identity spanning 81% of query; E-value: 4e-31) [73] and the kinase domain of *Arabidopsis* receptor kinase BRI1 (PDB: 5LPV; 30% identity spanning 66% of the query; E-value: 5e-27) [33]. The PBS1 model in S10 Fig was created much like the ZED1 model but using structures of BIK1 (PDB: 5TOS; 48% identity spanning 70% of query; E-value: 1e-97) [52] and BRI1 (PDB: 5LPV; 44% identity spanning 62% of query; E-value: 1e-71) [33] as model-building templates. Modeling was based on two templates so as to generate a model reliably representing the entire kinase domain.

For homology modeling of the hypothetical asymmetric ZED1-PBS1 heterodimer shown in S17 Fig, multiple candidate templates were considered (S18 Fig). ZRK-PBL dimers have been described as part of ZAR1-ZRK1-PBL2^UMP^ complexes observed by cryo-EM [17,18], however in these structures the electron density corresponding to PBL2^UMP^ is not well resolved (mean per-residue B-factors: ∼195, ∼225) and represents less than half of the PBL2 sequence (S18 Fig, panels A, B). Although a disordered, partly-unfolded state may be a biologically realistic consequence of PBL2 uridylation by the *X. campestris* effector AvrAC, the increased disorder of PBL2 relative to ZRK1 (and ZAR1) is also likely a result of the averaging of ZAR1-ZRK1-PBL2 particles - structural variation within and between particles increases in proportion to the radial distance from the centre of protein complexes [83]. While the structure of the PBL kinase BIK1 has been determined by X-ray crystallography [52] and is accordingly of higher resolution (mean per-residue B-factor: ∼57), in this case the kinase dimer interface is in a ‘back-to-back’ orientation that may be a crystal packing artefact and therefore not biologically-relevant (S18 Fig, panel C). In contrast, the X-ray crystal structure of human IRAK4 [42] is of similarly high resolution (mean per-residue B-factor: ∼64) and furthermore presents a ‘front-to-front’ interface in which the activation loops from both protomers are intimately involved in the dimer interface (S18 Fig, panel D). IRAK4 (PDB: 4U97) [42] was therefore used as model template for both ZED1 (30% identity across 63% of query sequence; E-value 7e-21) and PBS1 (42% identical across 62% of query sequence; E-value 8e-56) in addition to the top-scoring BLASTP hits for ZED1 (Pto) and PBS1 (BIK1) described above. In this case, however, activation loop residues from Pto and BIK1 were deleted from input alignments prior to modeling of ZED1 and PBS1 so as to force modeled activation loops to be constrained only by those from chains A and B from PDB entry 4U97 (IRAK4).

### Plant imaging and image processing

Photographs were acquired with a Nikon D5200 DSLR camera. We have previously described an ImageJ macro (PIDIQ, for <u>P</u>lant <u>I</u>mmunity and <u>D</u>isease <u>I</u>mage-based <u>Q</u>uantification) that quantifies pixel areas corresponding to green or yellow-shaded regions of input images [84]. Here we used a similar approach to replicate the behaviour of a factory-installed camera filter that serendipitously produces images with pixels representing dead/dying plant tissue and surrounding soil converted to grayscale, while those pixels representing healthy *Arabidopsis* leaf tissue remain unchanged. We determined that this behavior can be replicated by applying logic that outputs a full-colour, RGB pixel where the log-transformed ratio between pixel intensities for red and green channels 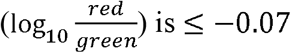, and a grayscale pixel where this condition is not satisfied (S20 Fig). Pixel analysis and image processing were implemented in PYTHON using MATPLOTLIB [68] and a script is available at https://github.com/DSGlab/Live-Dead-Filter.

## Author Contributions

DPB: Conceptualization, Investigation, Methodology, Validation, Formal Analysis, Software, Visualization, Writing – Original Draft Preparation, Writing – Review & Editing

MK: Investigation, Methodology, Resources, Writing – Review & Editing

AM, DS: Investigation, Methodology, Resources

IK: Investigation, Methodology

JZ: Methodology, Resources

WM, DM, JYL: Investigation

AHL: Methodology, Resources

YG: Data Curation

AS: Resources

DSG: Conceptualization, Data Curation, Funding Acquisition, Resources, Supervision, Writing – Original Draft Preparation, Writing – Review & Editing

DD: Conceptualization, Methodology, Funding Acquisition, Resources, Supervision, Writing – Original Draft Preparation, Writing – Review & Editing

## ACKNOWLEDGEMENTS

The authors wish to thank members of the Guttman and Desveaux labs for feedback and critical discussion of these data, with special thanks to Timothy Lo for discovery of the ‘Live/Dead’ camera filter. We are also grateful to Maël Baudin and Jennifer D. Lewis (University of California, Berkeley) for providing strains and advice for *ZAR1* gene-silencing, and to Jian-Min Zhou (Chinese Academy of Sciences, Beijing) for providing pUC19-35S-*avrAC*-HA. This work was supported by Natural Sciences and Engineering Research Council of Canada Discovery Grants (D.S.G and D.D.), Natural Sciences and Engineering Research Council of Canada Postgraduate Awards (D.S. and A.M.), Canada Research Chairs in Comparative Genomics (D.S.G.) and Plant-Microbe Systems Biology (D.D.), and the Center for the Analysis of Genome Evolution and Function (D.S.G. and D.D.).

## SUPPORTING INFORMATION

**S1 Fig. Yeast two-hybrid, three-hybrid and four-hybrid protein interaction schemes used to assess pairwise interactions between HopZ1a, ZED1/ZRKs, PBLs, and ZAR1.**

(A) Y2H: Transformants of haploid yeast strain EGY48 (*MAT* α) carrying pEG202 bait fusion plasmids (in this example, pEG202::*lexA^DBD^-ZED1*) are mated with transformants of haploid yeast strain RFY206 (*MAT* **A**) carrying pJG4-5 prey fusion plasmids (pJG4-5::*AD-PBL; AD = NLS-B42^AD^-HA*) and the reporter plasmid pSH18-34 (*lacZ*). (B) Y3H: Genes of interest (in this example *hopZ1a*) are integrated at the *ho* locus of the haploid yeast strain EGY48 (*MAT* α; see Materials and Methods). These strains were subsequently transformed with pEG202 bait fusion plasmids (in this example, pEG202::*lexA^DBD^-ZED1*) and mated with transformants of haploid yeast strain RFY206 (*MAT* **A**) carrying prey fusion plasmids (pJG4-5::*AD-PBL*) and the reporter plasmid pSH18-34. (C) Y4H: Transformants of haploid yeast strain EGY48 (*MAT* α) with integrated *hopZ1a* alleles and carrying pEG202 bait fusion plasmids (pEG202::*lexA^DBD^-ZAR1*) are mated with transformants of haploid yeast strain RFY206 (*MAT* **A**) with integrated *ZED1^wt^* and carrying prey fusion plasmids (pJG4-5::*AD-PBL*) and the reporter plasmid pSH18-34.

**S2 Fig. Pixel analysis to define a ‘relative interaction strength’ metric.**

(A) Averaged RGB pixel values for each of the yeast colonies presented in Fig 2, S4 Fig, S5 Fig, S8 Fig, and S9 Fig (see Materials and Methods) are plotted as a function of the log-transformed relative intensities of their blue (x-axis) and green (y-axis) channels. This plotting strategy results in a smooth arc describing the transition between strong (mostly blue) and weak (yellowish white) interactions, while spots representing array positions with no yeast colonies (agar plate only) form a distinct cluster below. The data were fitted to a quadratic function of the form *y* = *a*(*x* + *b*)^2^ + *cx* + *d* (plotted as a red line; *a* = −1.511, *b* = 0.67, *c* = 1.607, *d* = 0.456; R^2^ = 0.802), and the point at which this curve reaches its maximum 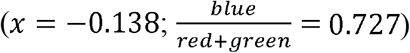 was used a threshold for discriminating strong interactions from bait-prey pairs with only 0.727 weak-to-moderate interaction strength. Spots exceeding this threshold are shown with a white border and are assigned white labels in the array layouts accompanying Fig 1A, Fig 2, Fig 5, S4 Fig, S5 Fig, S8 Fig, S9 Fig, and S11 Fig. (B) Histogram of the x-coordinates of all yeast spots plotted in panel A (excluding agar-only background spots). A consensus colour for the histogram bars representing each of 85 bins was determined by finding the average red, green, and blue channel intensities for all of the spots represented by a given bin. Bins exceeding the threshold described in panel A are highlighted with a white border. *Inset* - a colour-bar derived from the histogram data relating averaged pixel colours to a ‘relative interaction strength’ metric; the log-transformed relative intensities of the blue channel for each spot (−0.38 ≤ *x* ≤ 0.15) were scaled to values between 0 and 1. Empty bins (bins 76, 80, 81, 82, 83, and 84) were assigned the colour of the immediately-preceding non-zero bin, and the resulting colour range was smoothed by applying a five-bin sliding window average. The value corresponding to the white/black label threshold established in panel A (0.455) is indicated with a white horizontal line.

**S3 Fig. Peptides/modifications detected by LC-MS/MS at a more permissive threshold.** Peptides and acetylation sites from PBS1 (A) and ZED1 (B) are plotted as in Fig 1BC, except for the additional inclusion of peptides that only surpass the ‘Low’ confidence threshold reported by the Proteome Discoverer™ software, as described in S1 File.

**S4 Fig. AvrAC promotes interactions between ZRK1 and PBL kinases.**

*Top* - Binding of ZRK1 to 46 PBLs was assessed in the absence (Y2H; left) or presence (Y3H; right) of the *X. campestris* effector AvrAC; interaction schemes refer to S1 Fig. *Bottom* – Prey array layouts showing the relative strength of interactions between ZRK1 and PBLs, corresponding to the X-gal reporter plate above. Label colours (white or black) for each array position are determined by the relative interaction strength (see S2 Fig; Materials and Methods). Yeast spots and array positions corresponding to the positive control interaction between ZRK1 and PBL2 are highlighted with a red frame.

**S5 Fig. PBL-ZED1 interactions in the presence of HopZ2, HopZ3, and AvrAC.**

Y2H interactions between ZED1 and PBLs in the absence of effectors (panel B, left) are compared with Y3H interactions in the presence of the indicated effectors (panel A; panel B, middle). Prey array layouts at right indicate the relative strength of ZED-PBL interactions in the presence of HopZ2 (A) and in the absence of effector (B). All array labels are black since the relative interaction strengths are all below the threshold described in S2 Fig. Interaction schemes refer to S1 Fig. (C) *Top* - Representative western blots showing consistent expression of both bait (ZED1), prey (PBS1), and integrated effectors (HopZ alleles). *Bottom* – Nitrocellulose membranes stained with Ponceau S demonstrate equal loading of yeast cell extracts and consistent transfer from SDS-PAGE gels across all lanes.

**S6 Fig. Degenerate kinase motifs in ZED1 and the ZED1-related kinases (ZRKs).**

(A) Phylogenetic tree showing evolutionary relationships between the kinase domains of ZED1/ZRKs (dark blue shading), PBLs (light blue shading), and sequences from four structurally-characterized kinase domains - *Arabidopsis* BRI1 (PDB: 5LPV) [33] and BAK1 (PDB: 3UIM) [34], *Solanum pimpinellifolium* (tomato) Pto kinase (PDB: 3HGK) [73], and human IRAK4 (PDB: 4U97) [42]. Note that PBL28 is an out-group compared to other PBLs and may therefore not represent a ‘true’ PBL. (B) A subset of columns from the alignment used to generate the tree shown in panel A are presented to highlight important kinase motifs present in structurally-characterized kinase domains (top), ZED1/ZRKs (middle), and a representative subset of PBLs (bottom). The numbers of omitted, non-gap positions between adjacent blocks of consecutive columns are shown between square brackets for each sequence. Highlighted in blue are exact matches to established kinase motifs. Serine and threonine residues between the ‘DFG’ and ‘APE’ motifs that define the activation loop are highlighted in green. Serine/threonine residues that have been observed in phosphorylated form in previously-described crystal structures are highlighted in orange. PBS1 S244, a site acetylated by HopZ1a, and immediately following the site cleaved by HopAR1, is highlighted in red. PBL2 S253 and T254, sites uridylated by the *X. campestris* effector AvrAC and required for AvrAC-induced interaction with ZRK1 [12], are highlighted in purple. ZED1 residues targeted for mutagenesis in this study are indicated in bold, black font.

**S7 Fig. Active site residues in a homology-based model of pseudokinase ZED1 and in the experimentally-determined structure of receptor kinase BRI1.**

(A) Homology-based structural model (see Materials and Methods) of the ZED1 active site showing the relative positions of the sites targeted for mutagenesis in this study. Blue residue labels indicate positions that were mutated based on their established importance for kinase function. Red residue labels (and all-red sidechains) indicate positions acetylated by HopZ1a (S84, T87, T125, T177). The β3 lysine, K76 (labeled in grey), was not mutated but its sidechain is shown for comparison to the salt-bridge-forming lysine (K911) from BRI1 (shown in panel B, below). Note that in this model W58 and W193 would be expected to clash with the non-hydrolyzable ATP analogue AMP-PNP and Mn^2+^ ions if directly superimposed from the template BRI1 structure (shown in panel B). (B) An identical view of the kinase domain active site showing equivalent positions on the template kinase domain structure of the *Arabidopsis* brassinosteroid receptor kinase, BRI1 (PDB: 5LPV) [33]. Blue residue labels indicate positions aligned with those that were mutated in ZED1 based on their established importance for kinase function. Grey residue labels indicate positions equivalent to ZED1 sites that are acetylated by HopZ1a, as well as the salt-bridge forming β3 lysine, K911. (Note that Q919 and R922 were poorly resolved in the BRI1 structure, hence only the alpha-carbons of their sidechains are visible.) The ATP analogue AMP-PNP is shown as a ‘ball and stick’ representation, and two Mn^2+^ ions are shown as purple spheres. Images were prepared with UCSF CHIMERA [82].

**S8 Fig. Restoration of degenerate kinase motifs allows promiscuous, HopZ1a-independent ZED1-PBL interactions.**

(A) Protein interaction assays to assess the binding affinities of wild-type ZED1 and ZED1 single mutants (sites ‘b’, ‘c’, and ‘d’ in Fig 3) for 15 PBLs in the absence (Y2H; S1 Fig, panel A) or presence (Y3H; S1 Fig, panel B) of HopZ1a. The prey array layout at right indicates the relative strength of PBL interactions with ZED1^N173D^ in the absence of effector. The colours of array labels at each position (black or white) are determined by the relative interaction strength, as described in S2 Fig. (B) Double and triple mutant ZED1 alleles combining the mutations shown in panel A (sites ‘bc’, ‘bd’, ‘cd’, and ‘bcd’ in Fig 3) were likewise tested against the same array of 15 PBLs in both Y2H and Y3H contexts. The prey array layout shown at right indicates the relative strength of PBL interactions with ZED1^N173D W193G^ in the absence of effector. Interaction schemes refer to S1 Fig.

**S9 Fig. Restoration of a glycine-rich ZED1 ‘G-loop’ blocks HopZ1a-dependent and HopZ1a-independent PBL interactions.**

(A) Protein interaction assays to assess the binding affinities of ZED1^wt^, single mutants (ZED1^N173D^, ZED1^V212T^; sites ‘c’, ‘e’ in Fig 3), and a double mutant, (ZED1^“DT”^; sites ‘ce’ in Fig 3) for 15 PBLs in Y2H and Y3H contexts. The prey array layout shown at right indicates the relative strength of PBL interactions with ZED1^V212T^ in the absence of effector. The colours of array labels at each position (black or white) is determined by the relative interaction strength, as described in S2 Fig. (B) A ZED1 triple mutant (“3xG”; S56G W58G F61G) was tested against the same 15 PBLs, either alone or combined with the single and double mutants shown in panel A. The prey array layout shown at right indicates the relative strength of PBL interactions with ZED1^‘3xG’ V212T^ in the absence of effector. Interaction schemes refer to S1 Fig.

**S10 Fig. Glutamine substitutions of HopZ1a acetylation sites block ZED1-PBL interactions.**

(A) Structural models of ZED1 and PBS1 showing the positions of HopZ1a-acetylated residues as determined by LC-MS/MS. Note that acetylated PBS1 residues T32 and S405 do not form part of the conserved kinase domain. Images were prepared with UCSF CHIMERA [82]. (B) Y2H and Y3H assays for protein interactions between seven representative PBL preys and the indicated ZED1 bait alleles (wild-type, and mutants with glutamine substitutions at acetylation sites). (C) Y2H and Y3H assays for protein-protein interactions between wild-type ZED1 bait and the indicated PBS1 prey alleles (wild-type, and mutants with glutamine substitutions at acetylation sites). Interaction schemes refer to S1 Fig.

**S11 Fig. Isolated ZAR1 domains are not sufficient for robust formation of HopZ1a-dependent ternary complexes with ZED1 and PBLs.**

(A) Schematic showing ZAR1 domain truncation boundaries. Subdomains of the central nucleotide-binding region are labeled as described by Wang et al [17,18]: NBD, nucleotide-binding domain; HD1, helical domain 1; and WHD, winged helix domain. (B) Interactions between ZAR1 bait constructs and 11 PBL preys were assessed in the absence and presence of HopZ1a and/or ZED1 alleles integrated at the *ho* locus of strain EGY48 (*MAT* α) or RFY206 (*MAT* **A**), corresponding to Y2H (interaction scheme A), Y3H (interaction scheme B), and Y4H (interaction scheme C) assays (S1 Fig). The prey array layout shown at right indicates the relative strength of PBL interactions with ZAR1^LRR^ in the presence of both ZED1 and HopZ1a^wt^. All array labels are black since the relative interaction strengths are all below the threshold described in S2 Fig.

**S12 Fig. Dexamethasone-induced expression of ZED1^wt^ and ZED1^N173D^ alleles does not induce immunity in *Arabidopsis*.**

Control plants and BASTA-resistant *Arabidopsis* transformants bearing dexamethasone-inducible ZED1 alleles (ZED1^wt^ and ZED1^N173D^) were photographed before treatment (A) and 72 h post-dexamethasone spray (B) and screened for transgene expression (C), as described for ZED1^wt^ and ZED1^’DT’^ in Fig 6.

**S13 Fig. Dexamethasone-induced expression of ZED1^V212T^ causes whole-plant HR in Arabidopsis.**

Control plants and BASTA-resistant *Arabidopsis* transformants bearing dexamethasone-inducible ZED1^V212T^ were photographed before treatment (A) and 72 h post-dexamethasone spray (B) and screened for transgene expression (C), as described for ZED1^wt^ and ZED1^‘DT’^ in Fig 6.

**S14 Fig. BLASTP results showing sequences from *Nicotiana spp.* with similarity to *Arabidopsis* PBL and ZRK query sequences.**

Plotted points represent sequences from *Nicotiana spp.* (*N. tabacum*, *N. attenuata*, *N. sylvestris*, *N. tomentosiformis*) with regions homologous to the indicated *Arabidopsis* query sequences – 46 PBLs (blue) and 12 ZRKs (red). Up to 500 hits for each query are shown where the homologous region spans at least 50% of the length of the query sequence and has an expectation score (E-value) of less than 10e-20. Hits with E-values of zero were arbitrarily assigned values of 10e-225 prior to logarithmic transformation and plotting since the logarithm of zero is not defined.

**S15 Fig. Expression of a ZED1 allele with promiscuous PBL binding activity activates the Hypersensitive Response in *Nicotiana benthamiana*.**

(A) An *N. benthamiana* leaf infiltrated with *Agrobacterium* strains delivering the indicated (combinations of) transgenes, 26 h post-induction of expression by spray with dexamethasone. The image is representative of six different leaves infiltrated in the same way (two leaves each from three individual plants) and is consistent with multiple independent experiments. (B) Protein lysates from leaf cores punched from an additional leaf, similarly infiltrated with the same *Agrobacterium* cell suspensions shown in panel A, were assessed for expression of HA-tagged transgenes. (C) An *N. benthamiana* leaf infiltrated with *Agrobacterium* strains delivering the indicated (combinations of) transgenes, post dexamethasone spray, as for panel A. The image is representative of ten different leaves infiltrated in the same way (five replicates each in two independent experiments).

**S16 Fig. Survey of the prevalence of ambiguous and possible phospho/acetyl-accepting residues present in ZED1 sequences from diverse *Arabidopsis* ecotypes.**

(A) Histograms showing the incidence of ambiguous (left) or potential phospho/acetyl-accepting residues (right) in full-length ZED1 sequences (top) or in activation loop sequences only (bottom). (B) The same data shown in panel A are plotted as cumulative proportions of the total sample size. (C) Aligned ZED1 activation loop sequences for the Col-0 reference ecotype and the four *Arabidopsis* ecotypes with apparent phospho/acetyl-accepting residues. Matches to canonical DFG and APE motifs defining the activation loop are highlighted in blue, potential phospho/acetyl-accepting residues are highlighted in green, and V212 from the reference Col-0 ecotype is highlighted in bold, black font, as in S6 Fig, panel B. Ambiguous activation loop positions translated as ‘X’ are shown in bold, grey font.

**S17 Fig. Hypothetical ‘front-to-front’ pseudokinase-kinase heterodimer.**

(A) Molecular models of ZED1 (dark blue helices, orange activation loop) and PBS1 (light blue helices, pink activation loop) are superimposed over the asymmetric dimer of inactivated IRAK4 described by Ferrao *et al.* (PDB: 4U97) [42]. Note that the template IRAK4 structure is hidden except for the co-crystalized inhibitor, staurosporine, which competes with ATP for binding to kinase active sites. (B) A zoomed in view of the same hypothetical ZED1-PBS1 dimer shown in panel A, rotated by 90 degrees to highlight a ‘top-down’ view of the predicted interface between the ZED1 active site and the PBL activation loop acetylation site (S244 in PBS1). Images were prepared with UCSF CHIMERA [82].

**S18 Fig. Comparison of candidate templates for homology modeling of the ZRK-PBL pseudokinase-kinase interface.**

Experimentally-determined structures of kinase hetero- and homo-dimers (left column), and histograms of the average, per-residue B-factors for the two (pseudo)kinase protomers in each structure (middle and right columns, respectively). Kinase domains are depicted as ‘worm’ representations of the polypeptide backbone, with worm diameters proportional to the average B-factor at each position. As for S7 Fig, S10 Fig and S17 Fig, beta-strand elements are coloured yellow, while alpha-helices are coloured according to protein identity: dark blue (ZRK1, panels A, B); light blue (PBL2, panels A, B; BIK1, panel C); or brown (IRAK4, panel D). Activation loop sequences are coloured orange or pink. Uridylated side chains of PBL2 are shown in ‘ball-and-stick’ representation. Missing segments not observed in the structures are indicated with dashed lines. The proportion of the total sequence represented by a given structure is indicated above each histogram, expressed both as a fraction and as a percent. (A) The ZRK1-PBL2^UMP^ heterodimer from the monomeric nucleotide-free ZAR1-ZRK1-PBL2^UMP^ complex (PDB: 6J5V) [17]. (B) The ZRK1-PBL2^UMP^ heterodimer from the pentameric ZAR1-ZRK1-PBL2^UMP^ resistosome (PDB: 6J5T) [18]. (C) A homodimer of the *Arabidopsis* PBL kinase, BIK1 (PDB: 5TOS) [52]. (D) A homodimer of a catalytic site mutant (D311N) of the human kinase, IRAK4 (PDB: 4U97) [42].

**S19 Fig. Integration of catalytic segment and activation loop sequences with hydrophobic kinase domain ‘spines’ in ZAR1-ZRK1(-PBL2^UMP^) complexes.** ‘Catalytic’ and ‘regulatory’ spines (‘C-spine’, ‘R-spine’) for ZRK1 and PBL2^UMP^ are represented as molecular surfaces in brown (C-spines) and purple (R-spines). For clarity, ribbon representation of the protein sequences is restricted to three regions (where present in the structures): (1) β1-β2-β3-αC-β4-β5, (2) catalytic segment-activation loop, and (3) helix αF, which anchors both the catalytic and regulatory spines. The sidechain of ZRK1 residue M195 (equivalent to ZED1 acetylation site T177) is shown in red. The catalytic aspartate residues for ZRK1 (D191) and PBL2 (D219) are shown in green. Missing (unstructured) segments are indicated with dashed lines. C-spine and R-spine residues for ZRK1 and PBL2 are presented in S6 Table along with the corresponding positions from ZED1 and PKA [88]. Note that ZAR1 is also present in each of these structures (panels A-C) but is not shown for clarity. (A) ZRK1 in complex with ADP-bound ZAR1 - PDB entry 6J5W (B) ZRK1 (top) and PBL2^UMP^ (bottom) from a complex with nucleotide-free ZAR1 - PDB entry 6J5V. (C) ZRK1 (top) and PBL2^UMP^ (bottom) from a complex with ATP-bound ZAR1 - PDB entry 6J5T.

**S20 Fig. Demonstration of the pixel-processing logic used in transformations of *Arabidopsis* plant images.** Pixels from an input image (A) are subjected to threshold-masking of full-colour RGB pixels (C) based on the ratio between red and green channel intensities (B). False-colour representations of the absolute pixel intensities for individual red (D), green (E), and blue (F) channels are shown below.

**S1 Table. Plasmids generated and/or used for this study.**

Plasmids are organized by type into spreadsheet tabs: 1A – Gateway-compatible ‘entry vector’ clones; 1B – Y2H bait plasmids; 1C – Y2H prey plasmids; 1D – Yeast integration plasmids; 1E – Plant expression plasmids.

**S2 Table. Bait fusion plasmid transformants of yeast strain EGY48 (*MAT* α) and its derivatives bearing chromosomal integrations of bacterial effectors or *Arabidopsis ZED1*.**

**S3 Table. Prey fusion plasmid transformants of yeast strain RFY206/pSH18-34 (*MAT* A) and its derivatives bearing chromosomal integrations of *Arabidopsis ZED1*.**

**S4 Table. *Agrobacterium* strains used for transformation of *Arabidopsis* and *N. benthamiana*.**

**S5 Table. Oligonucleotide sequences used for site-directed mutagenesis.**

**S6 Table. Catalytic and regulatory spine residues for ZRK1, PBL2, ZED1, and PKA.**

**S1 File. Supplementary description of mass spectrometry analysis and results.**

**S2 File. Core kinase domain alignment used for phylogenetic inference of the trees shown in Fig 3B and S4 Fig, panel A.**

**S3 File. Newick format phylogenetic tree describing relationships between kinase domain sequences.**

**S4 File. CLUSTAL-formatted MUSCLE alignment of 813 unique ZED1 protein sequences representing 1121 distinct *Arabidopsis* ecotypes.**

**S5 File. List of *Arabidopsis* ecotypes corresponding to each of the unique ZED1 protein sequence IDs in S4 File.**

