## Supplemental File 1 for "Perturbations of the ZED1 Pseudokinase Activate Plant Immunity"

**S1 File – Supplementary description of mass spectrometry results**

**General comments**

LC-MS/MS analysis of trypsin-digested yeast lysates from cells co-expressing the *Arabidopsis* kinase PBS1 with wild-type (wt) or catalytic mutant (C216A) alleles of the *P. syringae* acetyltransferase effector HopZ1a were prepared in triplicate (three samples from independently-grown cultures for each of the two co-expressed HopZ1a alleles). Analysis of yeast lysates from cells co-expressing the *Arabidopsis* pseudokinase ZED1 with HopZ1a has been described previously [1], so here we have described a single sample each for cells co-expressing ZED1 with HopZ1a$^{wt}$ or HopZ1a$^{C216A}$, presented as controls to demonstrate consistency with these previous (and additional unpublished) results.

The peptide coverage/acetylation summaries presented in Fig 1BC and S3 Fig are derived from Microsoft Excel-formatted output generated by the Proteome Discoverer$^{TM}$ software application (Thermo Fisher Scientific), in which peptides (with and without post-translational modifications) are stratified according to the confidence level above which they are detected ('Low', 'Medium', and 'High'). The Low threshold results in detection of peptides representing 30.5-59.2%, 52.4-55.7% and 43.6-57.2% of the total sequences of PBS1, ZED1 and HopZ1a, respectively, while the represented proportions of the same three sequences range from 18.2-41.2%, 46.1-46.4% and 34.7-53.9% when evaluated with the Medium stringency threshold. Fig 1BC presents peptides/modifications that surpass High and/or Medium thresholds, while for contrast, S3 Fig presents the same samples shown in Fig 1BC but also includes peptides/modifications that only surpass the Low threshold. In addition to classification in this way (with proprietary software), all post-translational modifications described in the text were also independently identified and verified by manual inspection of chromatography peaks to identify peptides with mass increases of 42 Da (or multiples thereof) which are indicative of post-translational modifications that result in addition of (an) acetyl group(s). Selection of charged peptide ions from peaks of interest for fragmentation analysis, and subsequent inspection of the mass-shift profiles of their resultant b- and y-ion series together allowed identification of the specific peptide positions modified by acetylation, as described previously [2].

**HopZ1a acetylation sites on PBS1**

Based on the criteria described above, we identified four unique peptides indicating three distinct HopZ1a acetylation sites on PBS1. The PBS1 peptide 'Peptide 1' (KQSQP[T-Ac]VSNNISGLPSGGEK) was detected in all three of the samples co-expressing HopZ1a$^{wt}$ and none of the samples co-expressing HopZ1a$^{C216A}$. 'Peptide 2' (QSQP[T-Ac]VSNNISGLPSGGEK) was detected in just two of the three samples expressing HopZ1a$^{wt}$ and none of the samples co-expressing HopZ1a$^{C216A}$. It should be noted however that Peptides 1 and 2 are identical except for a single missed cleavage event and that both of these peptides indicate acetylation of PBS1 T32. 'Peptide 3' (LGPTGDK[S-Ac]HVSTR) was detected in all three of the samples co-expressing HopZ1a$^{wt}$. Although this peptide was not reliably detected with 'Medium' confidence in any of the three samples expressing HopZ1a$^{C216A}$ (Fig 1B), spectra consistent with this (unmodified) peptide were in fact observed using the more permissive 'Low' confidence threshold in two of these three samples (S3 Fig, panel A). We thus speculate that acetylation of the PBS1 activation loop alters its physicochemical properties so that it 'flies right' in the LC-MS/MS instrumentation whereas the unmodified peptide does not 'fly well'. Although this lower threshold increases the peptide coverage for all proteins across all samples and also identifies additional potential acetylation sites that do not meet our more stringent, manually-curated criteria (S3 Fig), we note that this relaxed (Low) threshold also results in the identification of acetylated S398, a modification that is detected in samples from cells co-expressing PBS1 with HopZ1a$^{C216A}$ as well (S3 Fig, panel A). Finally, 'Peptide 4' (NDDGGGSGSKFDLEG[S-Ac]EKEDSPR; representing acetylated S405) was observed in just one of three samples from cells co-expressing HopZ1a$^{wt}$ when subjected to the more stringent (Medium) threshold, but in all three of these samples when assessed at the Low stringency threshold. Importantly, none of the three PBS1 acetylation sites selected for mutagenesis in this study (T32, S244, S405) were modified in the presence of HopZ1a$^{C216A}$ - even at the lower-stringency threshold - suggesting that our methods and analysis are sensitive but also sufficiently conservative to guard against false positives that could possibly result from effector overexpression and/or promiscuous acetylation by endogenous yeast acetyltransferases.

**HopZ1a acetylation sites on ZED1**

Consistent with our previous report [1] we detected the ZED1 peptide, 'Peptide 5' (NINP[T-Ac]NIFIDENWTAK; representing acetylated T177) in lysates from cells co-expressing

*Arabidopsis* ZED1 with HopZ1a$^{wt}$ but not with HopZ1a$^{C216A}$ . In addition, we also detected modified ZED1 peptides ‘Peptide 6’ (WSSQNL[S-Ac]SFTEAYR) and ‘Peptide 7’ (WSSQNLSSF[T-Ac]EAYR) that represent acetylations of distinct residues from the same trypsin-generated peptide – S84 and T87, respectively (Fig 1C). While these two modifications are consistent with our additional unpublished results, acetylation of S137 (which is suggested by detection of ‘Peptide 8’; DGGL[S-Ac]SGVVLPWK; Fig 1C) is not; for this reason we did not pursue mutational analysis of S137. No additional acetylation sites were detected in either sample using the Low stringency threshold (S3 Fig, panel B).
